# Genetic variation reveals a homeotic long noncoding RNA that modulates human hematopoietic stem cells

**DOI:** 10.1101/2025.07.16.664824

**Authors:** Peng Lyu, Gaurav Agarwal, Chun-Jie Guo, Adam Sychla, Wallace Bourgeois, Tianyi Ye, Chen Weng, Mateusz Antoszewski, Samantha Joubran, Alexis Caulier, Michael Poeschla, Scott A. Armstrong, Silvi Rouskin, Vijay G. Sankaran

## Abstract

The *HOXA* gene locus coordinates body patterning, hematopoiesis, and differentiation. While studying blood phenotype-associated variation within the *HOXA* locus, we identified a genetic variant, rs17437411, associated with globally reduced blood counts, protection from blood cancers, and variation in anthropometric phenotypes. We find that this variant disrupts the activity of a previously unstudied antisense long non-coding RNA (lncRNA) located between *HOXA7* and *HOXA9*, which we have named *HOTSCRAMBL*. The *HOTSCRAMBL* variant disrupts lncRNA function and reduces human hematopoietic stem cell (HSC) self-renewal. Mechanistically, *HOTSCRAMBL* enables appropriate expression and splicing of *HOXA* genes in HSCs, most notably *HOXA9*, in an SRSF2-dependent manner. Given the critical role of *HOXA* gene expression in some blood cancers, we also demonstrate that *HOTSCRAMBL* variation or deletion compromises *HOXA*-dependent acute myeloid leukemias. Collectively, we show how insights from human genetic variation can uncover critical regulatory processes required for effective developmental gene expression.

## Introduction

The clustered homeobox (*HOX*) genes encode highly conserved transcription factors that are essential for embryonic development, particularly in establishing axial body patterning and defining segmental identity^1-4^. Among these, the *HOXA* locus, first discovered as regulators of body segmentation, have more recently been identified as critical for normal and malignant hematopoiesis^5-7^. For example, *HOXA* genes control the specification, self-renewal, and maintenance of human and mouse hematopoietic stem cells (HSCs)^8^. Moreover, several *HOXA* genes (especially *HOXA9*) are often overexpressed in acute myeloid leukemias (AMLs) as a result of activation by somatic driver events and predict a poor prognosis^9,10^. While these observations underscore the importance of tight regulation of the *HOXA* locus in hematopoiesis, the full spectrum of underlying regulatory mechanisms remains incompletely understood.

Several regulators of gene expression at the *HOXA* locus in HSCs have been identified. For example, *SOX17* has been shown to modulate *HOXA* gene expression in developmental hemogenic endothelium to initiate definitive hematopoiesis^11^. Moreover, direct binding of mixed-lineage leukemia (MLL) fusion proteins promote aberrant *HOXA* gene expression in leukemia^12^, and we have previously demonstrated a role for increased transcription elongation at the *HOXA* locus in HSCs driving inherited predisposition to developing myeloid malignancies^13^. Furthermore, three-dimensional chromatin interactions can maintain long-range enhancer-promoter interactions^14^. Disruption of a CTCF boundary between *HOXA7* and *HOXA9* has been shown to alter chromatin topology and activate aberrant posterior *HOXA* gene expression in AML, underscoring the critical role of genome architecture in regulating the *HOXA* locus^15^.

Long non-coding RNAs (lncRNAs) have also been shown to play critical roles in various aspects of *HOXA* gene regulation^16-18^. The lncRNA *Haunt* and its underlying genomic locus have been shown to exert opposing effects on *HOXA* gene activation during embryonic stem cell differentiation, highlighting the complexity of cis-regulatory control at this locus^19^. Importantly, antisense lncRNAs within the *HOXA* cluster itself exert direct regulatory effects on *HOXA* gene expression. For example, the lncRNA *HOTTIP* facilitates chromatin activation and looping across the *HOXA* genes^20^, and its overexpression enhances HSC self-renewal, contributing to AML pathogenesis^8^. *HOTAIRM1*, located between *HOXA1* and *HOXA2*, regulates *HOXA1* and *HOXA4* expression during myeloid differentiation, partly via splicing modulation^21^, and also shapes chromatin state and three-dimensional architecture to coordinate *HOXA* gene activation^22^. While these observations demonstrate a key role for lncRNAs and other regulators in modulating transcription at this locus, the role of human genetic variation on *HOXA* gene expression remains poorly characterized.

Here, while studying blood phenotype-associated variation at the *HOXA* locus, we made the serendipitous observation that there exists a poorly characterized lncRNA at the locus in the antisense direction between *HOXA7* and *HOXA9* that we found has a critical role in HSC self-renewal, which we termed *HOTSCRAMBL* for *<u>H</u>OXA* <u>O</u>ppositestrand <u>T</u>ranscript, <u>S</u>tem-<u>C</u>ell <u>R</u>egulator, <u>A</u>ntisense <u>M</u>id-cluster <u>B</u>etween <u>L</u>oci. Importantly, our studies of *HOTSCRAMBL* illuminate fine-tuning mechanisms for *HOX* gene regulation during development that have been recently-acquired in evolution.

## Results

### Genetic variation in a *HOXA* locus long non-coding RNA associated with altered hematopoiesis

Though genetic variation impacting hematopoiesis has been shown to alter *HOXA* gene expression indirectly, the contribution of genetic variation within this locus to hematopoiesis has not been previously studied. We therefore undertook a comprehensive assessment of germline variation associated with hematopoietic phenotypes at the locus. Through our analyses, we identified common variants across five independent haplotypes that were associated with various blood cell count phenotypes (**Figure 1A, Table S1**). Of these, the sentinel variant rs35355140 was associated with a global reduction in a range of measured blood counts, spanning white blood cells and various subtypes, platelets, and red blood cells - representative of all major hematopoietic lineages (**Figure S1A, Table S1**). Additionally, this variant was also associated with anthropometric phenotypes that might have contributions from *HOXA* gene expression, such as sitting height and reduced ankle spacing^23^ (**Figure S1A**). Importantly, this haplotype was further associated with reduced leukocyte telomere length (**Figure 1B**), suggesting it is associated with reduced HSC self-renewal^24,25^. Moreover, we observed reduced risk of myeloproliferative neoplasms (OR=0.89, 95% CI=0.82-0.95, *P*=0.001), a type of chronic blood cancer (with a greater effect seen with inclusion of *JAK2*-mutant clonal hematopoiesis as well), and loss of the Y (LOY) chromosome, a premalignant state^26,27^ (OR=0.84, 95% CI=0.81-0.87, p=7×10^-21^). We did not observe associations for other clonal hematopoietic phenotypes, although in some cases there might have been limited overall power to detect an association (**Figures 1C and S1B**). These observations suggest that this haplotype may modulate the function of the *HOXA* cluster in a primitive HSC compartment to resist age-related transformation of HSCs.

**Figure 1:**
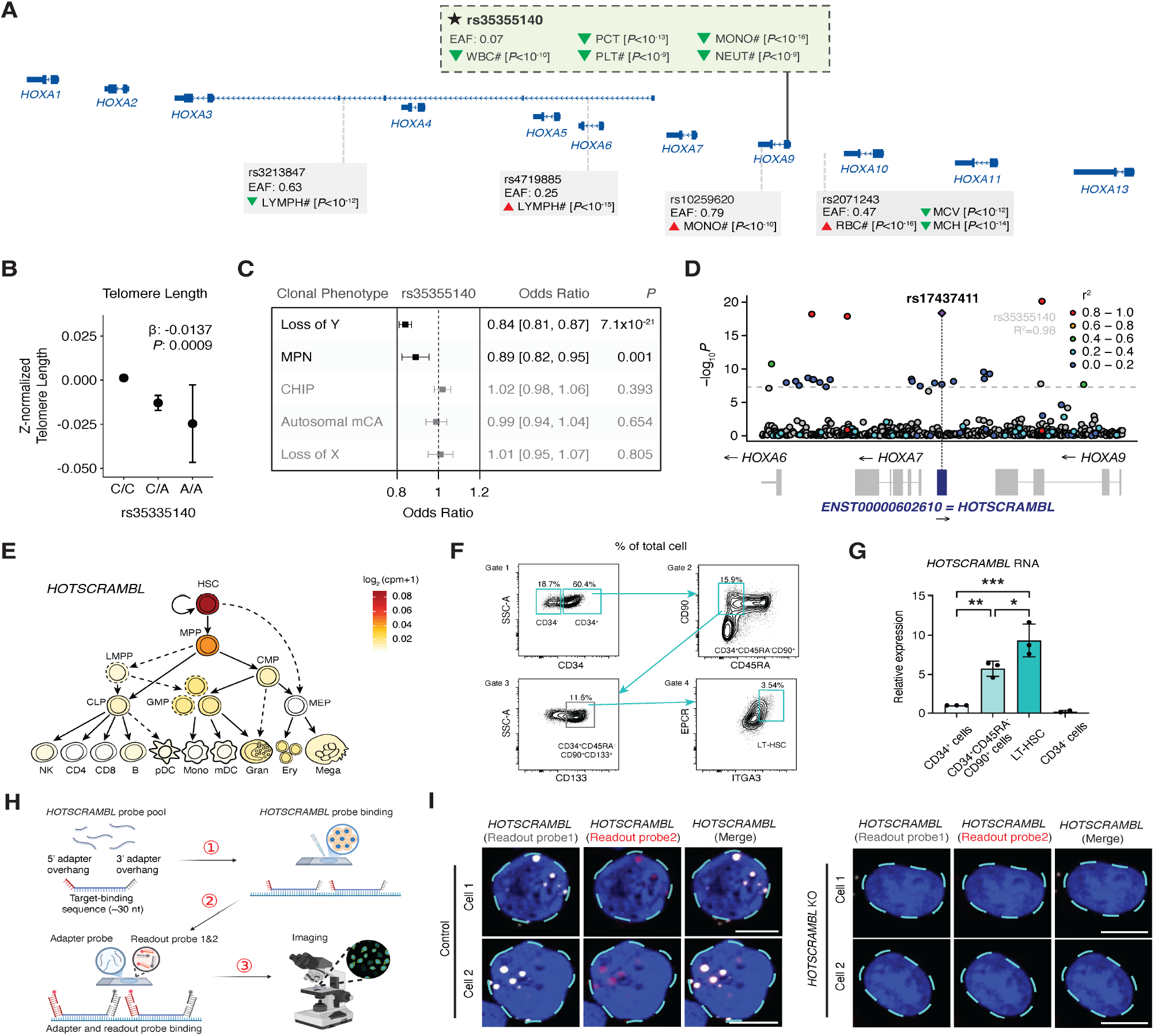
GWAS of blood cell phenotypes identifies rs17437411 within *HOTSCRAMBL*, a lncRNA highly enriched in human HSCs. **(A)** Genomic mapping of common variants at the *HOXA* locus associated with genome-wide significant alterations in blood cell traits, highlighting the sentinel variant rs35355140. EAF = effect allele frequency; LYMPH# = lymphocyte count; MONO# = monocyte count; RBC# = red blood cell count; MCV = mean corpuscular volume; MCH = mean corpuscular hemoglobin; WBC# = white blood cell count; NEUT# = neutrophil count; PLT# = platelet count; PCT = plateletcrit. **(B)** Z-normalized leukocyte telomere length by rs35335140 genotype in UK Biobank. Beta and p-values determined by linear regression analysis of the association between genotype dosage and z-normalized telomere length controlling for age, sex, and the first 10 genetic principal components. **(C)** Forest plot showing odds ratios and 95% confidence intervals for associations between rs35335140 and clonal hematopoietic phenotypes. MPN = myeloproliferative neoplasms; CHIP = clonal hematopoiesis of indeterminate potential; mCA = mosaic chromosomal alterations. **(D)** LocusZoom plot showing the loss of Y (LOY)-protective haplotype, identifying fine-mapped variant rs17437411 (R^2^=0.98 with sentinel variant rs35355140). **(E)** Expression of *HOTSCRAMBL* (log_2_ normalized counts per million mapped reads) throughout hematopoietic differentiation. **(F)** Representative flow cytometry gating strategy used to isolate HSPC subpopulations. Cyan boxes denote the sorted populations in (G). **(G)** Expression of *HOTSCRAMBL* in sorted human hematopoietic populations measured by quantitative PCR (qPCR). Long-term (LT)-HSC (CD34^+^CD45RA^−^CD90^+^CD133^+^EPCR^+^ITGA3^+^). **(H)** Schematic illustration of the MERFISH probe design and workflow used to detect *HOTSCRAMBL* transcripts. **(I)** Representative MERFISH images showing detection of *HOTSCRAMBL* transcripts in control and *HOTSCRAMBL* knockout cells. Nuclei (DAPI) are outlined by dashed cyan boundaries. Scale bars, 5 μm. All data are presented as mean ± standard deviations (SD), significance is indicated as \**P* < 0.05, \*\**P* < 0.01, or \*\*\**P* < 0.001.

We performed fine-mapping (**Figure 1D, Table S2**), identifying rs17437411 as the single likely functional variant in the haplotype. rs17437411 was predicted to potentially alter splicing or other activities of an intergenic 633 bp lncRNA (*ENST00000602610*) lacking an open reading frame (**Figure S1C**), located between *HOXA7* and *HOXA9. ENST00000602610* remains poorly characterized, and we hereafter term this lncRNA *HOTSCRAMBL* (<u>H</u>OXA <u>O</u>pposite-strand <u>T</u>ranscript, <u>S</u>tem-<u>C</u>ell <u>R</u>egulator, <u>A</u>ntisense <u>M</u>idcluster <u>B</u>etween <u>L</u>oci). Remarkably, the *HOTSCRAMBL* genomic DNA sequence is highly conserved among mammals (e.g., 94.5% in monkey, 71.2% in mouse), but shows low sequence similarity in non-mammalian vertebrates such as xenopus and zebrafish. In contrast, nearby *HOXA* genes, like *HOXA9*, exhibit strong cross-species conservation at the DNA coding sequence region (e.g., 97.9% in monkey, 93.1% in mouse, 70.7% in xenopus, and 59.2% in zebrafish) (**Figures S1D and S1E**). Moreover, no orthologs of this human lncRNA have been identified in other species.

Expression of *HOTSCRAMBL* within the hematopoietic system was largely restricted to CD34^+^ hematopoietic stem/progenitor cells (HSPCs), particularly within the most primitive HSCs (**Figures 1E-1G**). Indeed, while *HOTSCRAMBL*-positive cells were rare in bulk CD34^+^ HSPCs, CD34^+^CD45RA^-^CD90^+^ HSC-enriched populations demonstrated higher *HOTSCRAMBL* expression. Subcellular fractionation revealed predominant nuclear localization of *HOTSCRAMBL* (**Figure S1F**). Consistent with this, RNA FISH using two independent antisense probes detected distinct *HOTSCRAMBL* signals within the nucleus and at the nuclear periphery of CD34^+^CD45RA^−^CD90^+^ HSPCs (**Figures S1G and S1H**). In addition, Multiplexed Error-Robust Fluorescence *in situ* Hybridization (MERFISH) using independent readout probe sets targeting *HOTSCRAMBL* revealed a similar spatial distribution pattern, with discrete puncta detected within the nucleus and near the nuclear periphery (**Figures 1H and 1I**). While other lncRNAs within the locus, such as *HOTTIP* and *HOTAIR*, have been shown to play important roles in regulation of the *HOXA* cluster, the function of *HOTSCRAMBL* has not been characterized. Motivated by the observed hematopoietic associations of a variant within this lncRNA, we reasoned that further studies of this variant and *HOTSCRAMBL* could provide new functional insights.

### *HOTSCRAMBL*^rs17437411^ reduces long-term HSC maintenance *in vitro*

We sought to directly interrogate the role of *HOTSCRAMBL* and its associated rs17437411 variant in primary human CD34^+^ HSPCs (**Figure 2A**). To this end, we developed a high-fidelity CRISPR/Cas9-editing approach to model the effect of *HOTSCRAMBL*-KO (deletion of *HOTSCRAMBL*, without direct effects on other *HOXA* genes or regulatory elements) (**Figure S2A**), achieving ∼80% deletion efficiency (**Figures S2B and S2C**) and downregulating *HOTSCRAMBL* expression by ∼68% (**Figure S2D**). We also faithfully modeled the precise rs17437411 C>T variant in primary human CD34^+^ HSPCs using cytosine base editors (CBE), without introducing any undesired alterations to nearby bases (**Figure S2E**). We achieved editing at ∼75% of alleles in *HOTSCRAMBL* (**Figures S2F and S2G**), without affecting *HOTSCRAMBL* expression (**Figure S2H**). These edits were benchmarked against Cas9 or CBE-mediated editing at the *AAVS1* safe-harbor locus, respectively.

**Figure 2:**
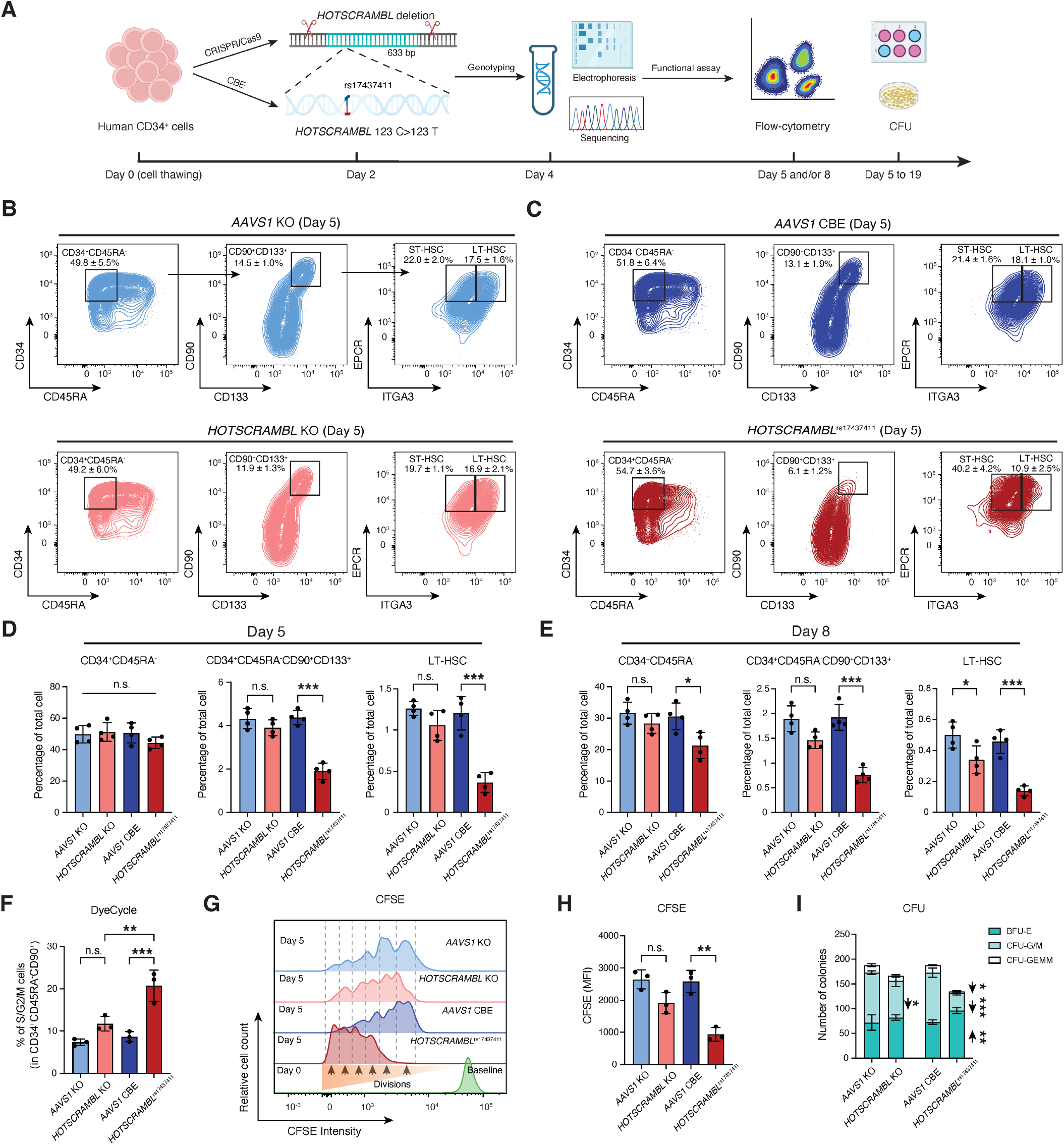
HOTSCRAMBLrs17437411 impairs human LT-HSC maintenance by promoting proliferation and differentiation in vitro. **(A)** Sche-matic overview of experimental design for CRISPR/Cas9 or cytosine base editing (CBE) mediated editing of *HOTSCRAMBL* at rs17437411 in human CD34^+^ HSPCs, followed by genotyping, RNA, and in vitro functional assays at indicated timepoints. **(B and C)** Representative flow cytometry plots of CD34^+^CD45RA^−^, CD34^+^CD45RA^−^CD90^+^CD133^+^, and LT-HSC (CD34^+^CD45RA^−^CD90^+^CD133^+^EPCR^+^ITGA3^+^) subsets at Day 5 post-editing in *AAVS1* KO vs. *HOTSCRAMBL* KO (B), and *AAVS1* CBE vs. *HOTSCRAMBL*^rs17437411^ (C). **(D and E)** Quantification of CD34^+^CD45RA^−^, CD34^+^CD45RA^−^CD90^+^CD133^+^, and LT-HSC populations at Day 5 (D) and Day 8 (E) post-editing based on flow cytometry. **(F)** Flow cytometric quantification of CD34^+^CD45RA^−^CD90^+^ cells in S/G2/M phases at Day 5 by DyeCycle staining. **(G)** Representative CFSE flow cytometry plots showing cell division profiles of CD34^+^CD45RA^−^CD90^+^ cells on Day 5. **(H)** Quantification of CFSE mean fluorescence intensity (MFI) in CD34^+^CD45RA^−^CD90^+^ cells on Day 5. **(I)** Stacked bar plots of colony-forming unit (CFU) assay at 14 days post-plating, quantifying the distribution of erythroid (BFU-E), granulo-cyte/macrophage (CFU-G/M), and multilineage granulocyte-erythrocyte-monocyte-megakaryocyte (CFU-GEMM) colonies. All data are presented as mean ± SD, significance is indicated as ∗*P* < 0.05, ∗∗*P* < 0.01, ∗∗∗*P* < 0.001, or n.s. (not significant).

Given that *HOTSCRAMBL* is selectively expressed in HSCs, we sought to assess its role in these cell compartments. While *HOTSCRAMBL*-KO mildly reduced the maintenance of phenotypic long-term repopulating (LT)-HSCs (CD34^+^CD45RA^-^CD90^+^CD133^+^EPCR^+^ITGA3^+^) in *in vitro* cultures of human CD34^+^ HSPCs (**Figures 2B and 2D**), *HOTSCRAMBL*^rs17437411^ editing resulted in a more profound decrease in LT-HSCs (**Figures 2C and 2E**). To uncover how the HSC population is lost after *HOTSCRAMBL*^rs17437411^ editing, we performed apoptosis and cell cycle assays (**Figure S2I**). While there was no increase in detectable apoptosis (**Figure S2J**), *HOTSCRAMBL*^rs17437411^ editing increased the proportion of CD34^+^CD45RA^-^CD90^+^ HSPCs in S/G2/M phase by 2.5-fold (**Figures 2F and S2K**), and increased their number of cell divisions 3-4 times more than the *AAVS1* CBE groups at 72 hours post-editing (**Figures 2G and 2H**).

These observations suggest that *HOTSCRAMBL*^*r*s17437411^ decreases both the maintenance and quiescence of primitive HSCs, which likely promotes differentiation of these cells, as we have observed for other perturbations previously^28^.

Consistent with this, when phenotypic LT-HSCs were enriched, we found that *HOTSCRAMBL*^rs17437411^ decreased the expression of canonical HSC marker genes, including *HLF, MECOM, CRHBP*, and *MLLT3* (**Figures S2L and S2M**). Furthermore, *HOTSCRAMBL*^rs17437411^ edited HSPCs showed a 7-fold loss of multipotent colony-forming unit (CFU) granulocyte, erythroid, macrophage, megakaryocyte (GEMM) colonies, and a reciprocal 1.5-fold increase in more differentiated burst-forming unit erythroid (BFU-E) colonies (**Figure 2I**). Taken together, these findings suggest that rs17437411 acts to reduce HSC maintenance and promote differentiation of these cells *in vitro*.

### *HOTSCRAMBL* loss-of-function promotes differentiation and reduces HSC self-renewal

To more rigorously assess how *HOTSCRAMBL* modulates human HSC function, we performed xenotransplantation of edited human CD34^+^ HSPCs into *Kit* mutant and immunodeficient NOD.Cg-*Kit*^*W-41J*^*Tyr*^+^*Prkdc*^*scid*^*Il2rg*^*tm1Wjl*^/ThomJ (NBSGW) mice (**Figure 3A**). Interestingly, *HOTSCRAMBL*^rs17437411^ edited cells initially demonstrated 3.8-fold greater peripheral chimerism at week 4 after transplantation, but 3-fold reduced chimerism once long-term engraftment was achieved at 16 weeks (**Figure 3B**). *HOTSCRAMBL*-KO also showed a 1.3-fold reduction in engraftment at 16 weeks post-transplantation (**Figure 3B**). Furthermore, while *HOTSCRAMBL*^rs17437411^ or KO edits were negatively selected against in bone marrow CD34^+^ HSPCs (**Figure 3C**), suggesting competitive disadvantage upon loss of *HOTSCRAMBL*, editing frequencies reciprocally increased in CD34-negative cells after transplantation (**Figure 3D**). Moreover, *HOTSCRAMBL* disruption resulted in increased commitment to early erythroid progenitors (CD71^+^CD235a^-^) in the bone marrow (**Figures S3A-S3C**). Taken together, these data are consistent with a failure of self-renewal and an increased propensity to differentiation upon perturbation or loss of *HOTSCRAMBL* in HSCs *in vivo*.

**Figure 3:**
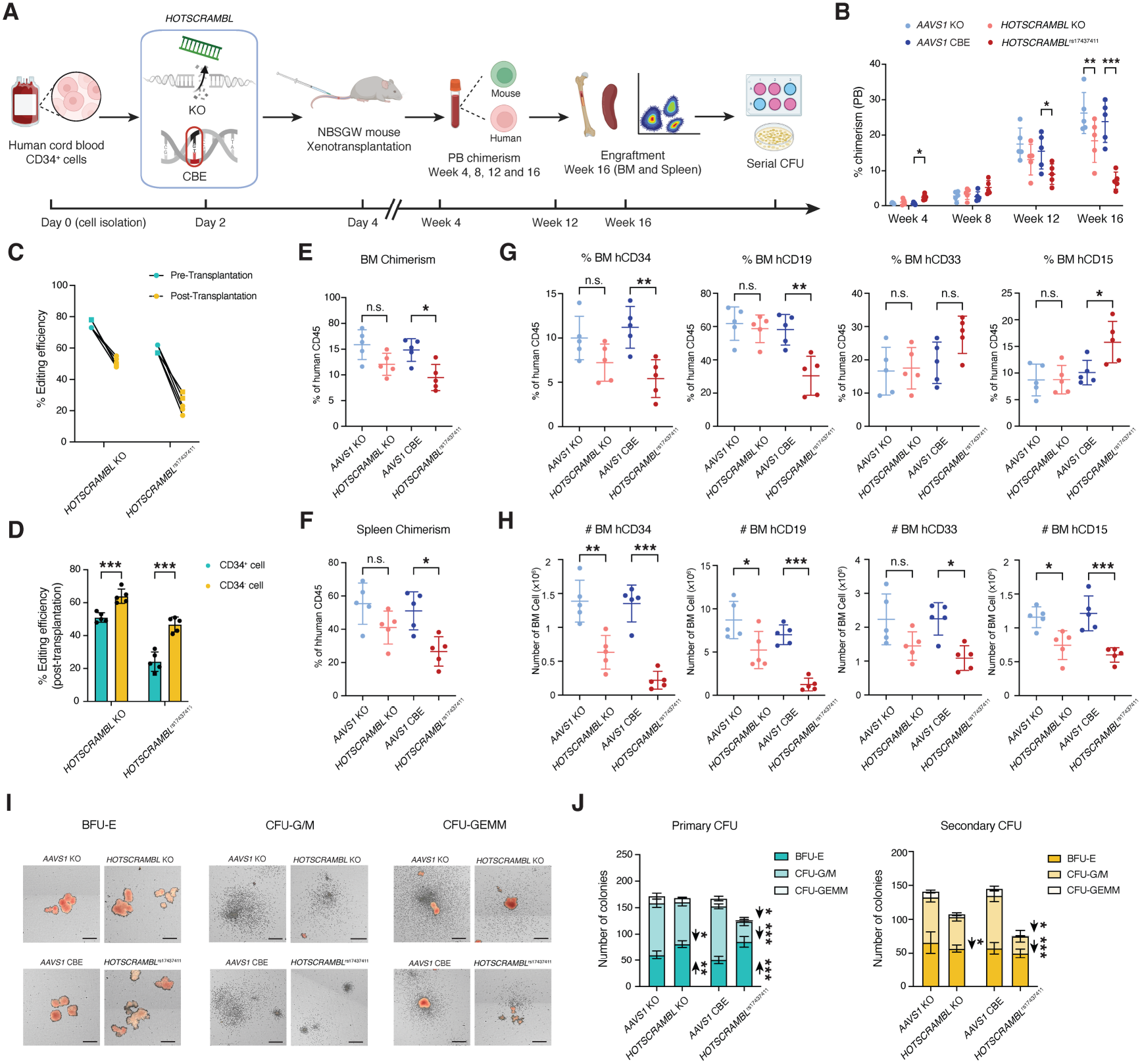
*HOTSCRAMBL*^rs17437411^ impairs long-term engraftment of human HSC and promotes differentiation *in vivo*. **(A)** Experimental work-flow for *HOTSCRAMBL* KO and CBE in human CB CD34^+^ cells, followed by xenotransplantation into NBSGW mice and longitudinal assessment of chimerism in peripheral blood and engraftment in bone marrow (BM) and spleen. **(B)** Human CD45^+^ PB chimerism (% human CD45^+^ of total human and mouse CD45^+^ cells) quantified by flow cytometry at 4, 8, 12, and 16 weeks post-transplantation. **(C)** Comparison of *HOTSCRAMBL* editing efficiency in input HSPCs before and after transplantation at week 16. **(D)** Comparison of *HOTSCRAMBL* editing efficiency in selected CD34^+^ and CD34^−^ human BM cells at week 16 post-transplantation. **(E and F)** Quantification of human CD45^+^ chimerism in BM (E) and spleen (F) at week 16 by flow cytometry. **(G)** Percentage of human BM blood lineage engraftment at week 16, including CD34^+^ HSPCs, CD19^+^ B cells, CD33^+^ myeloid progenitors, and CD15^+^ granulocytes by flow cytometry. **(H)** Number of human CD34^+^ HSPCs, CD19^+^ B cells, CD33^+^ myeloid progenitors, and CD15^+^ granulocytes in BM. **(I)** Representative images of BFU-E, CFU-G/M, and CFU-GEMM colonies from primary CFU assays. Scale bar, 200 µm. **(J)** Stacked bar plots of primary (left) and secondary (right) CFU assay at 10 days post-plating, quantifying the distribution of BFU-E, CFU-G/M, and CFU-GEMM colonies. All data are presented as mean ± SD, significance is indicated as ∗*P* < 0.05, ∗∗*P* < 0.01, ∗∗∗*P* < 0.001, or n.s. (not significant).

We next evaluated how *HOTSCRAMBL* regulates multilineage reconstitution of hematopoiesis (**Figure S3D**). Once long-term engraftment had been achieved, *HOTSCRAMBL*^rs17437411^-edited cells exhibited a significant reduction in human chimerism in both bone marrow (1.6-fold) and spleen (1.9-fold) (**Figures 3E and 3F**), without a significant reduction in overall human chimerism in *HOTSCRAMBL*-KO. Of engrafted human cells, *HOTSCRAMBL*^rs17437411^ editing reduced fractions of CD34^+^ HSPCs and CD19^+^ B-cells, but increased proportions of CD15^+^ granulocytes and monocytes. Nevertheless, *HOTSCRAMBL*-KO and *HOTSCRAMBL*^rs17437411^ editing both decreased absolute numbers of all lineages (**Figures 3G and 3H**). Moreover, *HOTSCRAMBL*^rs17437411^ edited CD34^+^ from the bone marrow resulted in reduced numbers and sizes of primitive CFU granulocyte/macrophage (CFU-G/M) and CFU-GEMM colonies upon colony replating (**Figures 3I, 3J, S3E, and S3F**). Importantly, there was a progressive loss of colony-forming potential with serial replating, suggesting that *HOTSCRAMBL*-editing reduces HSC self-renewal. Taken together, these data support the idea that rs17437411 impairs the self-renewal of human HSCs and promotes their differentiation, while deletion of *HOTSCRAMBL* has a similar, albeit milder, impact.

### The effect of rs17437411 on human HSCs is *HOTSCRAMBL*-dependent

While rs17437411 editing induced loss of human HSCs *in vitro* and *in vivo*, and milder phenotypes were seen with deletion of *HOTSCRAMBL* itself, we sought to validate that its effects were dependent on lncRNA activity (**Figure 4A**). *HOTSCRAMBL*^rs17437411^ edited cells (day 2) were targeted with two independent antisense locked nucleic acid (LNA) GapmeRs designed to specifically degrade *HOTSCRAMBL* and *HOTSCRAMBL*^rs17437411^ (**Figure S4A**). This strategy achieved >50% knock-down of both wild-type and *HOTSCRAMBL*^rs17437411^ edited transcripts in CD34^+^ HSPCs (**Figures S4B-S4D**). Strikingly, knockdown rescued the HSC loss following *HOTSCRAMBL*^rs17437411^ editing (**Figure 4B**), and partially restored colony-forming potential (**Figure 4C**), but showed minimal effect in controls, akin to the milder effects seen with *HOTSCRAMBL*-KO.

**Figure 4:**
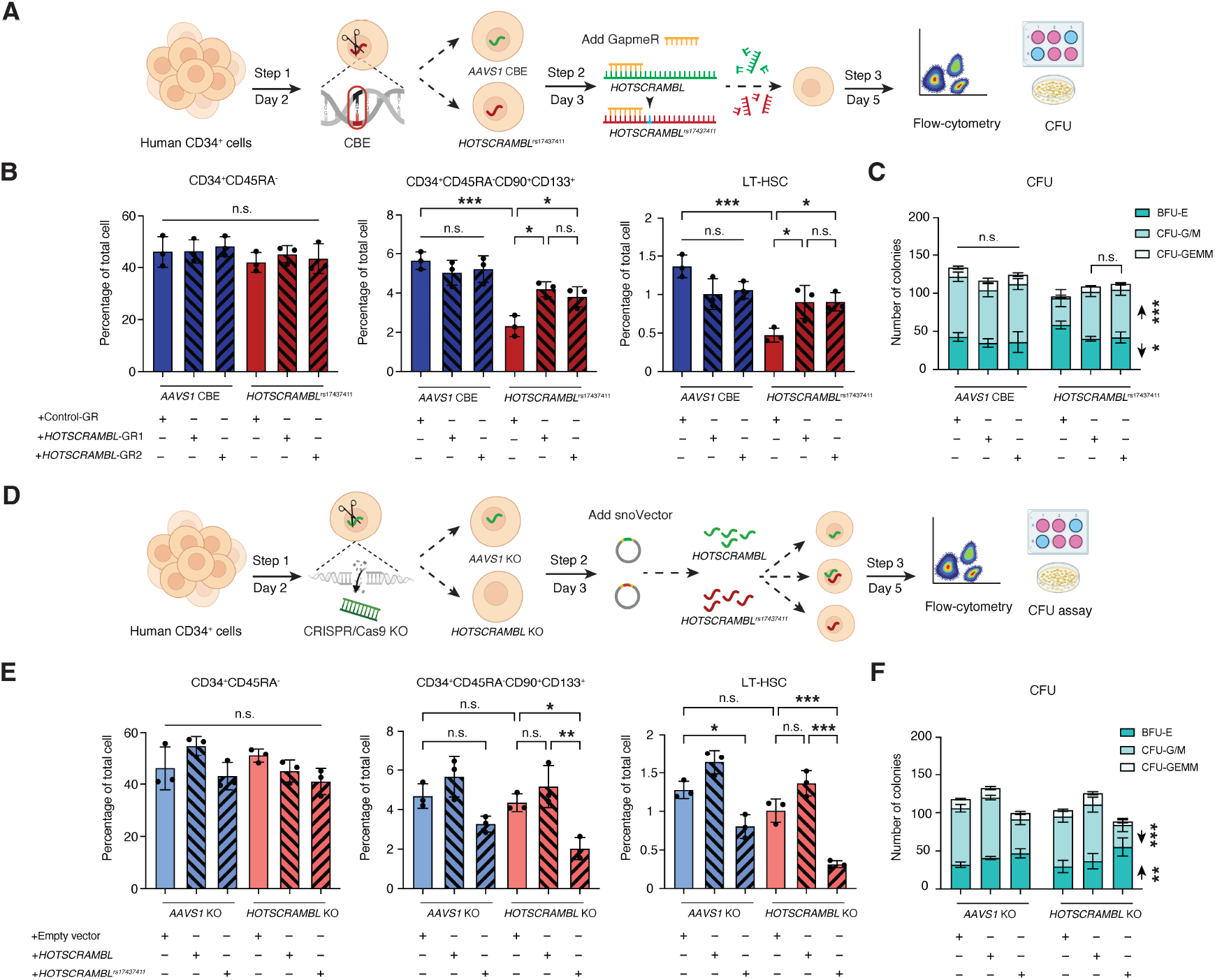
*HOTSCRAMBL*^rs17437411^ regulates LT-HSC maintenance in a transcript-dependent manner. **(A)** Experimental workflow for using Locked Nucleic Acid (LNA) GapmeR knockdown of HOTSCRAMBL in human CD34^+^ HSPCs (in *AAVS1* CBE and *HOTSCRAMBL*^rs17437411^). Cells were treated with HOTSCRAMBL-targeting or control GapmeRs and analyzed by flow cytometry and CFU assay at Day 5. **(B)** Flow cytometric quantification of CD34^+^CD45RA^−^, CD34^+^CD45RA^−^CD90^+^CD133^+^, and LT-HSC populations in cells treated with GapmeRs or control oligonucleotides under *AAVS1* CBE or *HOTSCRAMBL*^rs17437411^ editing conditions at day 5, respectively. GR = Gap-meR. **(C)** Stacked bar plots of CFU assay at 14 days post-plating, showing the distribution of BFU-E, CFU-G/M, and CFU-GEMM colonies. Cells were treated with GapmeRs or control oligonucleotides under *AAVS1* CBE or *HOTSCRAMBL*^rs17437411^ editing conditions, respectively. **(D)** Experimental workflow for overexpression of *HOTSCRAMBL* or *HOTSCRAMBL*^rs17437411^ RNAs following CRISPR/Cas9-mediated *HOTSCRAMBL* knockout in human CD34^+^ HSPCs. Cells were transduced with SnoVectors expressing *HOTSCRAMBL* or *HOTSCRAMBL*^rs17437411^, and assessed by flow cytometry and CFU at Day 5. **(E)** Flow cytometric quantification of CD34^+^CD45RA^−^, CD34^+^CD45RA^−^CD90^+^CD133^+^, and LT-HSC populations in cells overexpressed with *HOTSCRAMBL* or *HOTSCRAMBL*^rs17437411^ RNAs under *AAVS1* KO or *HOTSCRAMBL* KO editing conditions at day 5, respectively. **(F)** Stacked bar plots of CFU assay at 14 days post-plating, showing the distribution of BFU-E, CFU-G/M, and CFU-GEMM colonies. Cells were overexpressed with *HOTSCRAMBL* or *HOTSCRAMBL*^rs17437411^ RNAs under *AAVS1* KO or *HOTSCRAMBL* KO editing conditions at day 5, respectively. All data are presented as mean ± SD, significance is indicated as ∗P < 0.05, ∗∗P < 0.01, ∗∗∗P < 0.001, or n.s. (not significant).

We next tested whether expression of the *HOTSCRAMBL*^rs17437411^ RNA is sufficient to reduce HSC maintenance. Following *HOTSCRAMBL*-KO, we exogenously expressed *HOTSCRAMBL* in the nucleus using a SnoVector^29^ (**Figures 4D, S4E-S4G**). The exogenously expressed *HOTSCRAMBL* RNA localized predominantly in the nucleus (**Figure S4H**) and showed a ∼4.5-fold increase in expression over controls (**Figures S4I and S4J**). Following *HOTSCRAMBL*-KO, exogenous expression of *HOTSCRAMBL*^rs17437411^ transcript phenocopied the effects seen with endogenous genome editing to introduce rs17437411, resulting in a loss of LT-HSC maintenance (**Figure 4E**) and colony-plating potential (**Figure 4F**). Remarkably, expression of the mutant lncRNA could compromise the maintenance of LT-HSCs expressing wild-type *HOTSCRAMBL*, which suggests that *HOTSCRAMBL*^rs17437411^ may have dominant negative activity.

### *HOTSCRAMBL* modulates *HOXA9* expression in human HSCs

Having identified *HOTSCRAMBL* as a novel regulator of human HSCs, we sought to understand the underlying mechanisms. RNA-sequencing was performed in CD34^+^CD45RA^-^ CD90^+^ HSC-enriched cells following either *HOTSCRAMBL-* KO or *HOTSCRAMBL*^rs17437411^ editing (**Figure 5A**). While *HOTSCRAMBL*-KO edited cells exhibited only mild transcriptional changes, editing of the *HOTSCRAMBL*^rs17437411^ variant led to marked downregulation of several gene sets, including HSC signature genes, alongside upregulation of cell cycle-related genes, consistent with experimental observations (**Figures 5B, S5A, and S5B**). Notably, many *HOXA* genes, particularly *HOXA6, HOXA7*, and *HOXA9*, were significantly downregulated in the *HOTSCRAMBL*^rs17437411^ or KO groups, along with two antisense non-coding RNAs from that locus, *HOXA-AS2* and *HOXA-AS3* (**Figures 5B and 5C**). By more fully analyzing the observed gene expression changes, we found that while global genome-wide splicing event outcomes were similar across all groups (**Figure S5C**), KO or mutation of *HOTSCRAMBL* specifically altered splicing and/or expression of several *HOXA* genes, with the most drastic alterations at the highly expressed *HOXA9* mRNA (**Figures S5D and S5E**). This was interesting, in light of findings showing how lncRNAs transcribed near their target genes can modulate co-transcriptional splicing through direct interactions with chromatin or the splicing machinery^30– 32^. This might particularly make sense at the *HOXA* locus, where high-level transcriptional elongation is critical for altering expression of these genes in HSCs and other stem cell populations^13,33^.

**Figure 5:**
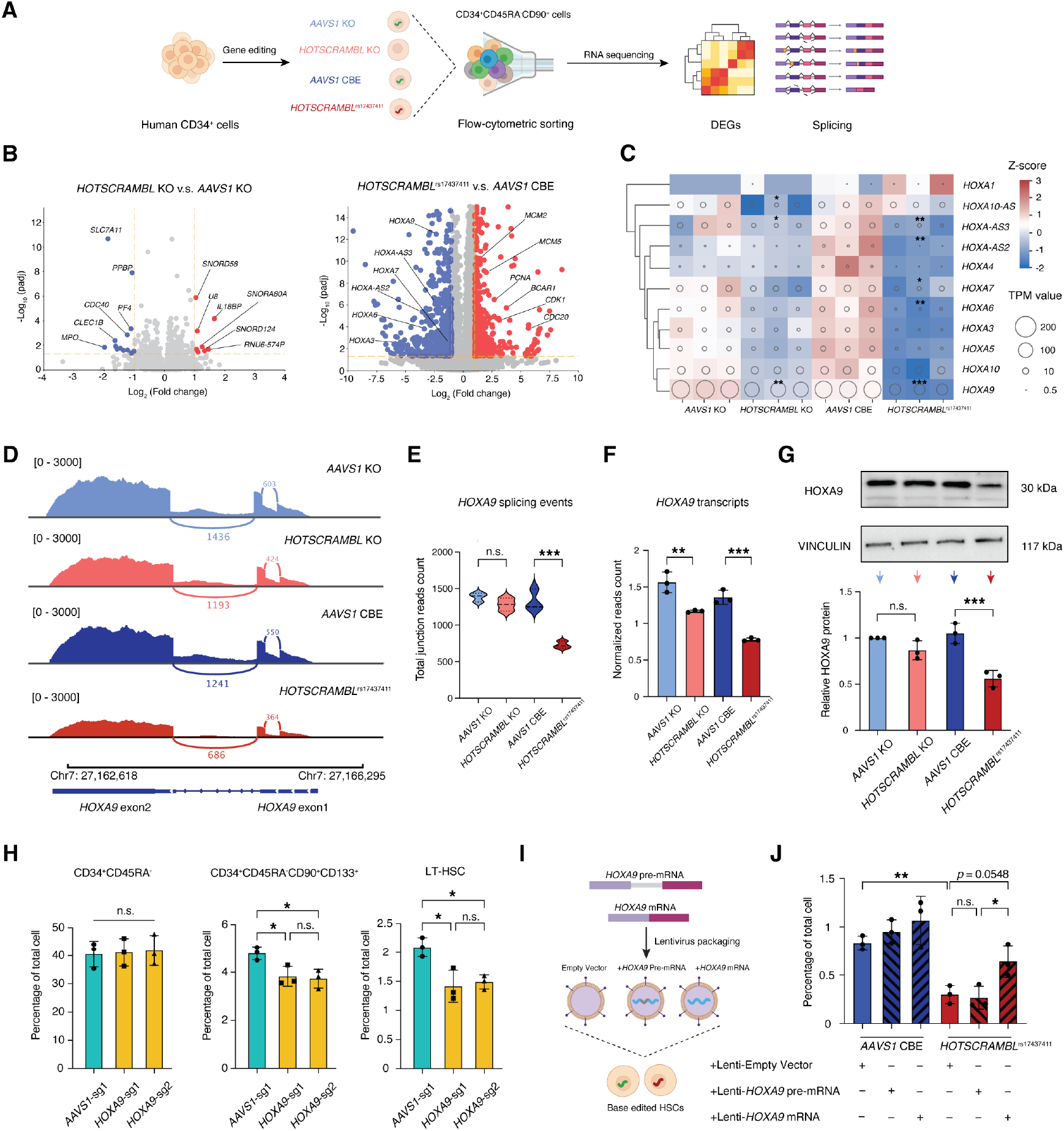
Reduced splicing efficiency underlies impaired HOXA9 expression upon HOTSCRAMBLrs17437411 editing. **(A)** Experimental workflow for gene editing, followed by enrichment of CD34^+^CD45RA^−^CD90^+^ HSPCs and bulk RNA sequencing. **(B)** Volcano plots showing DEGs in *HOTSCRAMBL* KO vs. *AAVS1* KO (left panel) and *HOTSCRAMBL*^rs17437411^ vs. *AAVS1* CBE (right panel). Values exceeding the plotting limits (|log2FC| > 10 or −log10(padj) > 15) are capped at the axis boundaries. **(C)** Heatmap of *HOXA* cluster genes in *AAVS1* KO, *HOTSCRAMBL* KO, *AAVS1* CBE and *HOTSCRAMBL*^rs17437411^ conditions. Colors indicate row-normalized TPM expression (z-score), and circle size indicates TPM value. Significance annotations were derived from DESeq2 analysis. **(D)** Sashimi plots of read coverage tracks at the *HOXA9* locus showing exon usage and splicing patterns. **(E)** Quantification of total splice junction read counts mapped to *HOXA9* exon 1-2. **(F)** Bar plot of normalized *HOXA9* transcript counts derived from mRNA sequencing. **(G)** Western blot analysis of *HOXA9* protein expression (up panel) and quantification normalized to VINCULIN (bottom panel). **(H)** Flow cytometric quantification of CD34^+^CD45RA^−^, CD34^+^CD45RA^−^CD90^+^CD133^+^, and LT-HSC populations in CD34^+^ HSPCs edited with *AAVS1*-sgRNA or two independent *HOXA9*-targeting sgRNAs (*HOXA9*-sg1 and *HOXA9*-sg2) using CRISPR/Cas9. **(I)** Schematic of lentiviral overexpression strategy using *HOXA9* pre-mRNA or mature mRNA in *AAVS1* CBE or *HOTSCRAMBL*^rs17437411^ edited groups, respectively. **(J)** Rescue of LT-HSC frequency in *AAVS1* CBE or *HOTSCRAMBL*^rs17437411^ edited group by lentiviral expression of *HOXA9* pre-mRNA or mRNA, quantified by flow cytometry. All data are presented as mean ± SD, significance is indicated as \**P* < 0.05, \*\**P* < 0.01, \*\*\**P* < 0.001, or n.s. (not significant).

*HOXA9* was particularly notable among the most differentially spliced genes following *HOTSCRAMBL*^rs17437411^ editing, given the magnitude of the change. *HOTSCRAMBL*^rs17437411^ editing impaired productive splicing at the intronic junction between exon 1 and exon 2 of *HOXA9* (**Figures 5D and 5E**) and resulted in reduced levels of *HOXA9* mRNA (**Figure 5F**), consistent with this change being a direct effect resulting from *HOTSCRAMBL*^rs17437411^. Importantly, we validated that *HOTSCRAMBL*^rs17437411^editing further reduced HOXA9 protein expression in CD34^+^CD45RA^-^ CD90^+^ HSC-enriched cells (**Figure 5G**), and recapitulated a *HOXA9* knockdown signature (**Figure S5A**).

Together, these data suggested that *HOTSCRAMBL* may regulate the expression of *HOXA9* and other genes at the locus. This was notable, given that *HOXA9* is highly enriched in human HSCs, is a known potent regulator of primitive HSC states, and that its overexpression is associated with exceptionally poor prognosis in acute myeloid leukemia (AML).

We next validated how alteration of *HOXA* genes, especially *HOXA6, HOXA7* and *HOXA9*, which were impacted by *HOTSCRAMBL* perturbation, could affect human HSCs. We performed gene knockout experiments with *HOXA6, HOXA7*, and *HOXA9* targeting guide RNAs. All guide RNAs showed ∼80% editing efficiency (**Figure S5F**). Interestingly, we found that *HOXA9* perturbation reduced LT-HSC maintenance *in vitro*, while perturbation of *HOXA6* or *HOXA7* alone did not (**Figures 5H and S5G**), suggesting that reduced *HOXA9* levels may largely contribute to the HSC loss observed following *HOTSCRAMBL*-editing. To further confirm the functionality of differential *HOXA9* splicing, we performed *HOTSCRAMBL*^rs17437411^ editing and sought to rescue HSC loss through lentiviral expression of either *HOXA9* pre-mRNA (containing the intron) or spliced mRNA (**Figure 5I**). Importantly, only expression of the spliced *HOXA9* mRNA was sufficient to partially rescue the loss of LT-HSCs (**Figure 5J**), demonstrating that *HOTSCRAMBL*^rs17437411^ functionally modulates human HSCs through reduced *HOXA9* splicing and expression. showed ∼80% editing efficiency (**Supp Fig. 5F**). Interestingly, we found that *HOXA9* perturbation reduced LT-HSC maintenance *in vitro*, while perturbation of *HOXA6* or *HOXA7* alone did not (**Fig. 5H and Supp Fig. 5G**), suggesting that reduced *HOXA9* levels may largely contribute to the HSC loss observed following *HOTSCRAMBL*-editing. To further confirm the functionality of differential *HOXA9* splicing, we performed *HOTSCRAMBL*^rs17437411^ editing and sought to rescue HSC loss through lentiviral expression of either *HOXA9* pre-mRNA (containing the intron) or spliced mRNA (**Fig. 5I**). Importantly, only expression of the spliced *HOXA9* mRNA was sufficient to partially rescue the loss of LT-HSCs (**Fig. 5J**), demonstrating that *HOTSCRAMBL*^rs17437411^ functionally modulates human HSCs through reduced *HOXA9* splicing and expression.

### *HOTSCRAMBL* orchestrates SRSF2-dependent splicing of *HOXA9*

We next wondered how rs17437411 and deletion of *HOTSCRAMBL* may functionally impair *HOXA9* splicing. We first predicted that the rs17437411 variant might alter the secondary structure of *HOTSCRAMBL* based on *in silico* modeling, which indicated local conformational differences between the wild-type *HOTSCRAMBL* and *HOTSCRAMBL*^rs17437411^ sequences. To experimentally validate this, we performed dimethyl sulfate mutational profiling with sequencing (DMS-MaPseq) on *in vitro*-transcribed *HOTSCRAMBL* and *HOTSCRAMBL*^rs17437411^ RNA^34^. We found that *HOTSCRAMBL*^rs17437411^ gains an alternative structure that is not present in the wild-type *HOTSCRAMBL*. While the wild-type *HOTSCRAMBL* primarily adopts a single stable conformation, the *HOTSCRAMBL*^rs17437411^ gives rise to two alternative structural states (α and β) with fractional representations of 0.56 and 0.44, respectively (**Figure S6A**). Rolling correlation of DMS signal and structural modeling showed that the *HOTSCRAMBL*^rs17437411^-α conformation displayed minor differences compared with the wild type, whereas the *HOTSCRAMBL*^rs17437411^-β conformation exhibited substantial structural rearrangements (**Figures S6B and S6C**). These data indicate that rs17437411 substantially alters the structural equilibrium of *HOTSCRAMBL*, which may in turn affect its RNA-RNA or RNA-protein interactions, and thereby interfere with downstream function.

To understand the direct mechanism through which *HOTSCRAMBL* alters *HOXA9* splicing, we sought to identify binding targets of *HOTSCRAMBL* (**Figure 6A**). We conducted these analyses in two AML cell lines (MOLM13 and OCI-AML4) that displayed robust *HOTSCRAMBL* and *HOXA9* expression at comparable levels to human CD34^+^ HSPCs at both the RNA and protein levels (**Figures S6D-S6G**). Following *HOTSCRAMBL*-KO or HOTSCRAMBL^rs17437411^ editing (**Figures S6H and S6I**), *HOTSCRAMBL* transcripts were targeted with several biotinylated probes and pulled down, and bound mRNAs were identified through Chromatin Isolation by RNA Purification followed by sequencing (ChIRP-seq)^35^. While *HOTSCRAMBL*-KO abolished RNA pulldown (**Figure S6J**), validating our approach, *HOTSCRAMBL*^rs17437411^ altered binding to select transcripts, including *HOXA9. HOTSCRAMBL* showed enriched association with intronic and exon-intron boundary regions of *HOXA9* (**Figure 6B**) and the rs17437411 variant reduced *HOTSCRAMBL* occupancy across the *HOXA9* gene body by approximately ∼2-fold (**Figure 6C**), with evident loss across the spliced intron, which we further validated through ChIRP-qPCR (**Figure 6D**).

**Figure 6:**
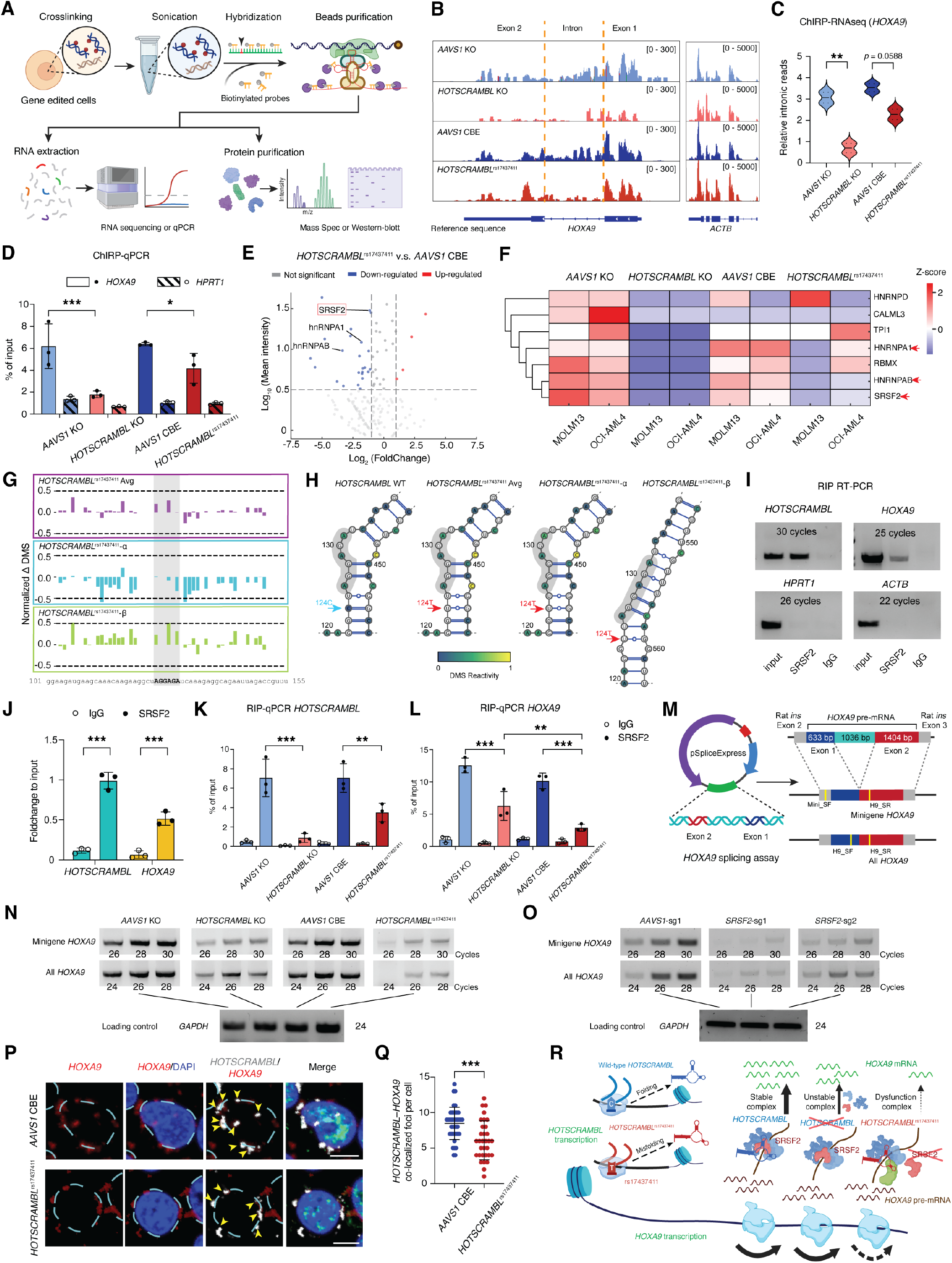
*HOTSCRAMBL*^rs17437411^ impairs *HOXA9* expression by weakening SRSF2-mediated pre-mRNA splicing. **(A)** Schematic workflow of comprehensive identification of RNA-binding proteins by mass spectrometry (ChIRP-MS) and chromatin isolation by RNA purification (ChIRP) followed RNA sequencing. **(B)** Sashimi plot showing normalized ChIRP-RNA sequencing read coverage at the *HOXA9* locus (left) and *ACTB* locus (right). **(C)** Quantification of normalized *HOXA9* intronic reads from (B). **(D)** Quantification of *HOXA9* in purified *HOTSCRAMBL*-interacting RNA fractions by chromatin isolation by RNA purification (ChIRP)-qPCR. *HPRT1* (control). **(E)** Volcano plot showing proteins identified by ChIRP-MS comparing *HOTSCRAMBL*^rs17437411^ vs. *AAVS1* CBE. SRSF2 was highlighted in red box. **(F)** Heatmap of ChIRP-MS data showing normalized abundance of selected splicing factors and RNA-binding proteins interacting with *HOTSCRAMBL* or *HOTSCRAMBL*^rs17437411^. Heat bar, normalized Z-score. **(G)** Normalized change in DMS signal (*HOTSCRAMBL* WT minus [*HOTSCRAMBL*^rs17437411^-Avg (purple peaks), *HOTSCRAMBL*^rs17437411^-α (cyan peaks) and *HOTSCRAMBL*^rs17437411^-β (green peaks)]) around the 126 SRSF2 binding site (grey shaded area). **(H)** Predicted RNA structure at the SRSF2 126-binding site colored by DMS reactivity. **(I)** RIP RT-PCR validating *HOTSCRAMBL* and *HOXA9* enrichment in SRSF2-pulldown fractions compared to IgG control. *ACTB* and *HPRT1* serve as negative controls. **(J)** Relative quantification of RT-PCR band intensities from (I), analyzed using ImageJ 8-bit grayscale measurement. **(K)** RIP-qPCR quantification of *HOTSCRAMBL* enrichment in SRSF2-IP compared to IgG control. **(L)** RIP-qPCR quantification of *HOXA9* enrichment in SRSF2-IP compared to IgG control. **(M)** Schematic of *HOXA9* minigene construct (pSpliceExpress^56^) used for splicing assay in MOLM13. Minigene-derived transcripts were detected using Mini-SF and H9-SR primers, while total *HOXA9* was detected using H9-SF and H9-SR. **(N)** RT-PCR analysis of minigene and all *HOXA9* splicing in *AAVS1* KO, *HOTSCRAMBL* KO, *AAVS1* CBE, and *HOTSCRAMBL*^rs17437411^ edited MOLM13 cells. *GAPDH* as loading control. **(O)** RT-PCR analysis of minigene and all *HOXA9* in MOLM13 cells edited with *SRSF2*-sg1, *SRSF2*-sg2, or *AAVS1* control sgRNA. *GAPDH* as loading control. **(P)** Representative RNA FISH images showing colocalization of *HOTSCRAMBL* (white) and *HOXA9* pre-mRNA (red) in *AAVS1* CBE control and *HOTSCRAMBL*^rs17437411^-edited CD34^+^ HSPCs. Nuclei (DAPI) are outlined with dashed blue circles. Co-localization of *HOTSCRAMBL* and *HOXA9* is indicated by yellow arrows. Scale bar, 5 μm. **(Q)** Quantification of *HOTSCRAMBL*-*HOXA9* co-localization per cell as determined by fluorescence imaging. **(R)** Schematic model illustrating that *HOTSCRAMBL* facilitates SRSF2 recruitment to *HOXA9* nascent transcripts, promoting co-transcriptional splicing and efficient *HOXA9* mRNA production. All data are presented as mean ± SD, significance is indicated as \**P* < 0.05, \*\**P* < 0.01, \*\*\**P* < 0.001, or n.s. (not significant).

To further define underlying mechanisms, we performed comprehensive identification of RNA-binding proteins by mass spectrometry (ChIRP-ms)^36^. Interestingly, a set of proteins under 50 kDa showed decreased binding in the *HOTSCRAMBL*^rs17437411^ edited group (**Figure S6K**). Moreover, from the ChIRP-MS, 41 and 80 directly bound proteins were identified in MOLM13 and OCI-AML4 cell lines, respectively (Table S7). Strikingly, *HOTSCRAMBL*^rs17437411^ editing reduced binding of *HOTSCRAMBL* to several splicing factors, including members of the hnRNP complex, and to SRSF2 by ∼2-fold in terms of quantified peptides (**Figures 6E and 6F**). Given that DMS-MaPseq revealed marked structural rearrangements in *HOTSCRAMBL*, we next asked whether these conformational changes could abrogate SRSF2 interaction. Examination of DMS reactivity at predicted SRSF2 binding motifs showed reduced accessibility at two sites, one of which was located near the variant at nucleotide 126 and another at nucleotide 308^37^ (**Figures 6G and S6L**). Although the SRSF2 binding sites remained exposed in the wild-type and *HOTSCRAMBL*^rs17437411^ structures, DMS-constrained structural modeling of the alternative conformation indicated that these motifs became partially occluded (**Figures 6H and S6M**). This local closing of the RNA structure in the *HOTSCRAMBL*^rs17437411^ alternative form likely underlies the loss of SRSF2 association observed above. Consistent with this model, RNA immunoprecipitation (RIP)-qPCR showed that SRSF2 can directly bind to both *HOTSCRAMBL* and *HOXA9* mRNA (**Figures 6I and 6J**). Remarkably, *HOTSCRAMBL*^rs17437411^ editing was sufficient to abrogate ∼70-80% of SRSF2 binding to both *HOTSCRAMBL* and *HOXA9* (**Figures 6K and 6L**), suggesting that efficient SRSF2 interaction with *HOXA9* mRNA or pre-mRNA is dependent on *HOTSCRAMBL*.

To further define the mechanistic relationship between *HOTSCRAMBL, HOXA9*, and SRSF2 in primary CD34^+^ HSPCs, we performed combined RNA FISH and immunofluorescence in *HOTSCRAMBL*^rs17437411^ or control edited cells. These analyses revealed robust co-localization of *HOTSCRAMBL-*containing puncta with both *HOXA9* pre-mRNA and SRSF2 nuclear foci, with a remarkable reduction in *HOXA9* pre-mRNA expression in *HOTSCRAMBL*^rs17437411^-edited cells (**Figures S6N and S6O**). To evaluate the relative stoichiometry of *HOTSCRAMBL* and *HOXA9* pre-mRNA, we performed absolute quantification with qPCR in MOLM13 cells with control *AAVS1* CBE or *HOTSCRAMBL*^rs17437411^ editing. While HOTSCRAMBL RNA levels remained unchanged between groups, *HOXA9* pre-mRNA copy number per cell was significantly reduced in *HOTSCRAMBL*^rs17437411^-edited cells, and the ratios of *HOXA9* pre-mRNA to *HOTSCRAMBL* were approximately 2.6 in *AAVS1* CBE and 1.7 in *HOTSCRAMBL*^rs17437411^ samples, respectively (**Figure S6P**), indicating a selective reduction of *HOXA9* expression in *HOTSCRAMBL*^rs17437411^-edited cells. To test direct requirements for splicing, we performed a mini-gene splicing assay, in which the minimal *HOXA9* exon-intron-exon junction was expressed in MOLM13 cells, and efficient splicing of *HOXA9* was assessed via RT-PCR (**Figure 6M**). *HOTSCRAMBL*-KO and *HOTSCRAMBL*^*rs17437411*^ editing was sufficient to decrease *HOXA9* splicing of both mini-gene and endogenous *HOXA9* transcripts (**Figure 6N**), validating the role of *HOTSCRAMBL* in promoting efficient splicing of this transcript. Strikingly, SRSF2 perturbation with guide RNAs was sufficient to drastically reduce splicing efficiency of both the mini-gene construct and endogenous *HOXA9* (**Figures 6O and S6Q**). Consistently, high-resolution RNA FISH further revealed marked co-localization of *HOTSCRAMBL* with *HOXA9* pre-mRNA, supporting its role in co-transcriptional splicing. Notably, this co-localization was reduced in *HOTSCRAMBL*^rs17437411^-edited cells, reinforcing the functional disruption caused by the variant (**Figures 6P and 6Q**). Taken together, these data suggest that *HOTSCRAMBL* helps orchestrate SRSF2 localization to *HOXA9* to promote efficient splicing. The rs17437411 variant likely disrupts this interaction by altering *HOTSCRAMBL* secondary structure, thereby decoupling RNA-RNA and RNA-protein interaction and reducing *HOXA9* mRNA production (**Figure 6R**).

### Genetic modulation of *HOTSCRAMBL* induces differentiation in human AML

In human AML, high expression of *HOXA9*, as well as that of *HOXA6, HOXA7*, and the non-coding genes *HOXA-AS2* and *HOXA-AS3*, is individually associated with high-risk disease and poor clinical outcomes (**Figures S7A and S7B**). Given our functional data demonstrating that *HOTSCRAMBL* regulates efficient splicing of *HOXA9* and expression of other *HOXA* genes in human HSCs, we interrogated the relevance of *HOTSCRAMBL* expression in human AML. We observed that high *HOTSCRAMBL* expression portends a worse prognosis in both childhood and adult AML (**Figure 7A**).

**Figure 7:**
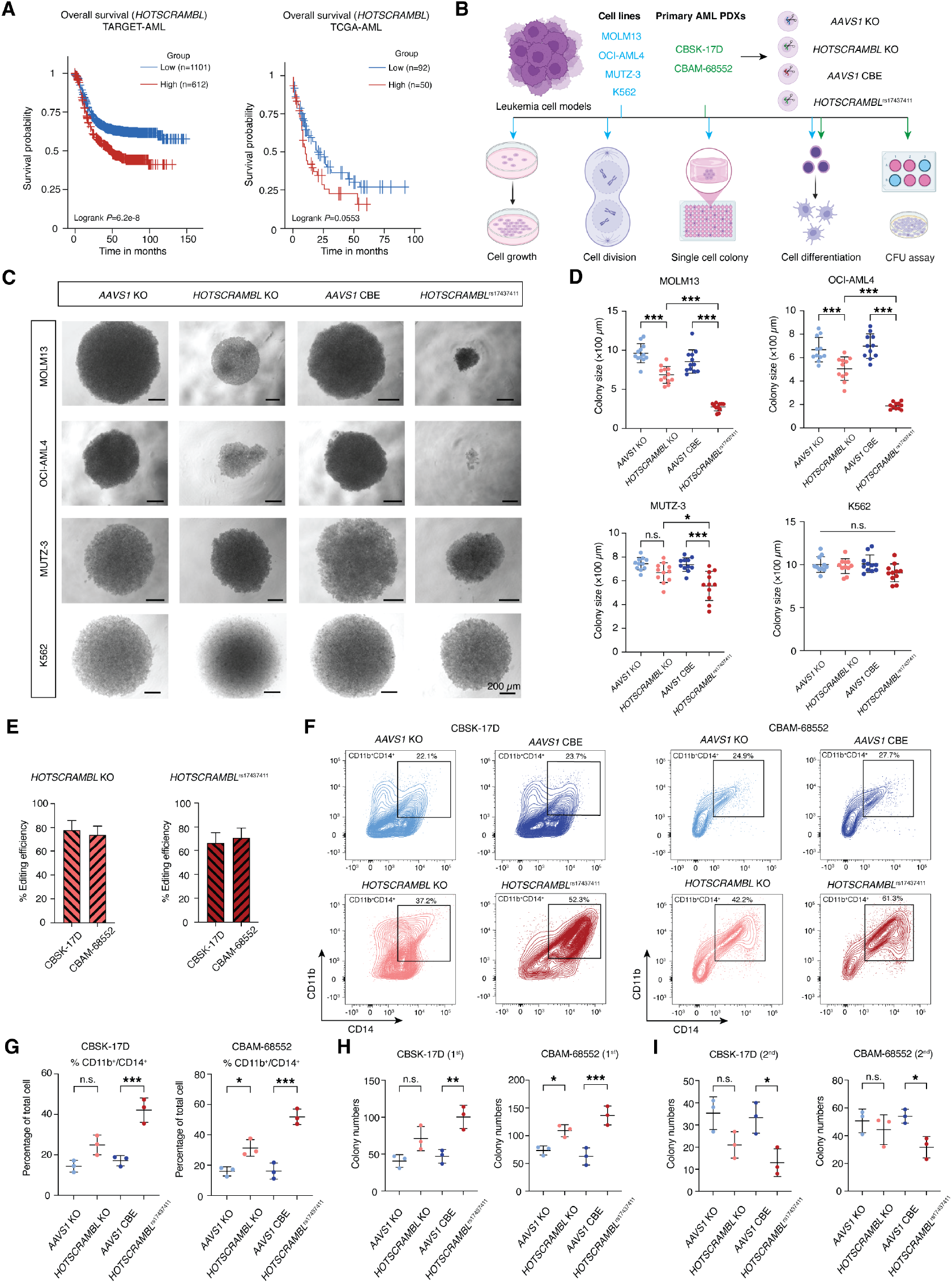
*HOTSCRAMBL*^rs17437411^ impairs leukemic cell growth and colony formation in *HOXA9*-dependent leukemia cell models. **(A)** Kaplan-Meier survival curve from the TARGET-AML and TCGA-AML dataset generated using the Survival Genie V2 platform^57^. **(B)** Experimental workflow assessing the impact of *HOTSCRAMBL* KO and *HOTSCRAMBL*^rs17437411^ edited groups across different leukemia cell lines and primary AML patient-derived cells. **(C)** Representative bright-field images of colonies formed from single cells at Day 14 post-plating in MOLM13, OCI-AML4, MUTZ-3, and K562 cells. Scale bar, 200 μm. **(D)** Quantification of colony sizes from (C). Each dot represents an individual colony. **(E)** Quantification of *HOTSCRAMBL* KO and base editing efficiency in CBSK-17D and CBAM-68552 at Day 4 post-editing. **(F)** Representative flow cytometric plots of the percentage of CD11b^+^CD14^+^ cells in CBSK-17D and CBAM-68552 at Day 4 post-editing. **(G)** Flow cytometric quantification of the percentage of CD11b^+^CD14^+^ cells in CBSK-17D and CBAM-68552 at Day 4 post-editing. **(H and I)** Primary (1^st^) and secondary (2^nd^) CFU assays performed using CBSK-17D and CBAM-68552. Cells were cultured in methylcellulose for 14 days per round, and total colonies were quantified. All data are presented as mean ± SD, significance is indicated as \**P* < 0.05, \*\**P* < 0.01, \*\*\**P* < 0.001, or n.s. (not significant).

We next examined how altered expression or function (rs17437411) of *HOTSCRAMBL* may perturb *HOXA*-dependent AMLs. We performed genetic perturbations in a variety of human AML cell lines (MOLM13, OCI-AML4, and MUTZ-3), all of which display robust expression of both *HOTSCRAMBL* and *HOXA9* (**Figure 7B**). As a negative control, we also performed editing in K562 cells, which have only minimal endogenous expression of *HOTSCRAMBL* and *HOXA9. HOTSCRAMBL*^rs17437411^ editing reduced *HOXA9* expression, as observed in human HSCs (**Figure S7C**), and decreased cell growth by ∼2-fold in MOLM13 and OCI-AML4 cells, and ∼1.5-fold in MUTZ-3 cells, respectively (**Figure S7D**). While K562 cells showed no change in cell cycle progression, *HOTSCRAMBL*^rs17437411^ editing markedly reduced proliferation in MOLM13 and OCI-AML4 cells. Conversely, in MUTZ-3 cells, which are more stem cell-like and contain a high proportion of CD34^+^ HSPCs^28^, editing led to increased cell cycle progression, suggestive of a differentiation phenotype similar to that observed in human HSCs (**Figure S7E**). Furthermore, there was reduced single-cell-derived colony growth following *HOTSCRAMBL*^rs17437411^ editing of MOLM13, OCI-AML4, and MUTZ-3 cells (**Figures 7C and 7D**), showing how *HOTSCRAMBL* perturbations can compromise clonogenic potential in a range of genetically and phenotypically diverse *HOXA*-dependent AML cell lines. A hallmark of aggressive AMLs is the maintenance of a more primitive undifferentiated population among a heterogenous group of cells^38^. We performed immunopheno-typing to understand how *HOTSCRAMBL* editing altered cellular composition in AML cell lines. K562 cells showed no changes in the composition of phenotyped populations following *HOTSCRAMBL*^rs17437411^ editing (**Figure S7F**). In contrast, *HOTSCRAMBL*^rs17437411^ editing increased the proportions of mature, differentiated CD11b^+^CD14^+^ and CD14^high^ myeloid populations in MOLM13, and CD11b^+^ granulocytic and CD14^+^ monocytic populations in OCI-AML4 cells (**Figures S7G-S7J**). Additionally, it was associated with decreased CD34^+^ progenitor-like populations and increased differentiated CD33^+^ myeloid populations in MUTZ-3 cells (**Figures S7K and S7L**).

To further validate these findings, we examined the effects of *HOTSCRAMBL* perturbation in primary AML cells obtained from patient-derived xenografts. Editing efficiency for both *HOTSCRAMBL* KO and *HOTSCRAMBL*^rs17437411^ was comparable across two patient samples (**Figure 7E**). Flow cytometry analysis showed that while control groups maintained baseline differentiation, both *HOTSCRAMBL* KO and particularly *HOTSCRAMBL*^rs17437411^ led to an increase in differentiated CD11b^+^CD14^+^ myeloid populations (**Figures 7F and 7G**). In colony forming assays, *HOTSCRAMBL* KO and *HOTSCRAMBL*^rs17437411^ cells exhibited increased total colony numbers compared to controls upon primary plating (**Figure 7H**). However, upon secondary replating, colony numbers markedly declined, suggestive of a depletion of self-renewing progenitor populations (**Figure 7I**). These data suggest that *HOTSCRAMBL* not only modulates *HOXA* gene expression in the setting of healthy human hematopoietic stem cells, but is also required in leukemias that co-opt many of the homeotic gene programs required to enable stem-cell like self-renewal properties.

## Discussion

The *HOXA* genes are critical regulators of normal HSC selfrenewal and are frequently dysregulated to drive aggressive myeloid malignancies, including high-risk AML^15,39^. Despite their established importance in human disease, much of what is currently known about the regulation of the *HOXA* locus has been inferred from studies in model organisms^40^. Several lncRNAs in the *HOXA* locus have been characterized to play roles in both health and disease. *HOTAIR* has been shown to reprogram chromatin states to promote metastatic breast cancer^41^. *HOXA-AS2* can enable epigenetic suppression of HBV circular DNA transcription^42^. *HOXA-AS3* and its paralog *HOXB-AS3*, two conserved lncRNAs at the center of the *HOXA* cluster, act in *cis* to regulate nearby *HOX* genes during early development^43^. *HOXA10-AS* has been identified as an oncogenic lncRNA in gliomas, regulating key developmental and cancer-related pathways^44^. Leveraging the unique vantage point provided by a naturally-occurring human genetic variant, here we have functionally characterized a previously unstudied lncRNA, which we term *HOTSCRAMBL*. We show that *HOTSCRAMBL* is critical to enable effective splicing and expression of *HOXA* genes in tandem with SRSF2 and other splicing factors, which alters HSC self-renewal and likely impacts other body patterning phenotypes.

Interestingly, the *HOTSCRAMBL* rs17437411 variant produced stronger phenotypic effects than complete deletion of the lncRNA, suggesting a possible dominant-negative mechanism. The structural change introduced by this variant appears to alter RNA conformation and impair SRSF2 binding, allowing the mutant transcript to compete for or sequester essential splicing or transcription factors. This interpretation is supported by our DMS-MaPseq data showing that *HOTSCRAMBL*^rs17437411^ adopts an alternative structure that occludes the SRSF2 binding interface. Nevertheless, we cannot exclude a potential gain-of-function or neomorphic component, as the altered RNA structure may also create novel interactions or stabilize non-canonical RNP complexes not formed by the wild-type lncRNA. Mutations in the splicing factor SRSF2, particularly the P95H mutation, alter exon recognition in a sequence-specific manner, driving recurrent mis-splicing of key hematopoietic regulators and contributing to impaired hematopoiesis and myelodysplasia^44–47^. These mutations can affect the way SRSF2 functions, leading to aberrant splicing and contributing to the development of blood cancers. We show that *HOTSCRAMBL* orchestrates the effective recruitment of the splicing factor SRSF2 to the *HOXA* genes, such as *HOXA9*. Whether and how *HOTSCRAMBL* activity connects to SRSF2 oncogenic mutations will be an important area to investigate in future studies. Additionally, perturbations of *HOXA6* or *HOXA7* individually cause slight reductions in LT-HSC maintenance, suggesting that the rs17437411 variant may affect multiple *HOXA* genes beyond *HOXA9*, either via an impact on splicing or expression. Coordinated downregulation of *HOXA* genes supports a broader *cis*-regulatory role of *HOTSCRAMBL*, similar to other *HOXA*-associated lncRNAs such as *HOTTIP*^*48*^ and *HOTAIRM1*^*22*^.

While such lncRNAs are generally detected only at low levels and in select cell types, our study showcases the value of human genetic variation to prioritize their function, and shows how HSC-enriched expression enables *HOTSCRAMBL* to have potent and cell-type specific function, at least within the hematopoietic compartment. Moreover, given that such antisense transcripts are ubiquitously annotated across the genome, it is likely that lncRNA-mediated splicing regulation may be an important layer of regulatory control at other genetic loci.

## Conclusion

Our study reveals *HOTSCRAMBL* as a regulator of human HSC function. Genetic variation predisposing to developing mosaic LOY and other clonal disorders has been shown to significantly increase the risk for acquiring blood cancers, highlighting the relevance of inherited variants in modulating HSC dynamics to promote premalignant and malignant states^26,49^. The rs17437411 variant is associated with a range of phenotypes in the UK Biobank, including a decreased risk of LOY, a lower risk of developing myeloproliferative neoplasms and related disorders, subtle reductions in a diverse set of blood cell counts, and shorter leukocyte telomere length. All of these phenotypes are consistent with reduced HSC self-renewal, which would limit the opportunity for clonal expansions to arise in the HSC compartment. Moreover, we show experimentally that *HOTSCRAMBL*^rs17437411^ editing or KO restrains human AML cells. Given that *HOTSCRAMBL*^rs17437411^ suppresses AML growth and promotes differentiation, it likely represents a blood cancer resilience variant, reminiscent of a distinct recently identified variant in the *MSI2* enhancer^47,50^.

From a broader perspective, our study motivated through the discovery of a phenotype-associated genetic variant illustrates how, although *HOX* gene regulation has been known to be highly conserved across evolution^4,51^, modulatory strategies that alter *HOX* gene expression have likely been coopted in the course of more recent evolution to fine-tune this process, such as we characterize for the intricate splicing and expression regulation mediated via *HOTSCRAMBL*.

## Limitations of the study

Our study has focused on hematopoietic phenotypes due to the rs17437411 variant in *HOTSCRAMBL*. However, it is likely given other associated phenotypes, that this lncRNA also modulates *HOXA* gene expression in other contexts (such as skeletal development). Further studies building from our observations here will likely extend our knowledge of this fine-tuned regulation across tissues. Moreover, while we illuminate the extent of hematopoietic plasticity by variable HSC self-renewal due to a naturally-occurring genetic variant, the extent to which such plasticity might be compensated for in the bone marrow of individuals, such as through progenitor expansion, remains to be studied by examining *in vivo* human hematopoietic phenotypes^52,53^. Finally, we have focused on the role of SRSF2 in mediating splicing phenotypes in cooperation with *HOTSCRAMBL*, but it is likely that other regulators of splicing, such as hnRNPs, are also involved. Deeper, low-input mechanistic validation in primary HSPCs would be valuable. This is an important area for future investigation.

## Supporting information

Source Data for All Figures

## Acknowledgements

We thank members of the Sankaran lab and other colleagues for valuable advice and guidance. We thank the Thermo Fisher Center for Multiplexed Proteomics at Harvard Medical School for assistance with mass spectrometry analysis. We thank Pu Zhen for guidance on MERFISH and for kindly sharing probes. This work was supported by the Howard Hughes Medical Institute (to V.G.S.), National Institutes of Health grants R01DK103794, R01HL146500, R01CA265726, R01CA292941 (to V.G.S.), the Gates Foundation (to V.G.S.), the Mathers Foundation (to V.G.S.), the Edward P. Evans Foundation (to V.G.S.), Blood Cancer United (to V.G.S.), the Alex’s Lemonade Stand Foundation (to V.G.S.), and a gift in memory of Jan Ellen Paradise, MD to Boston Children’s Hospital (to V.G.S.). Schematics in figures were created using BioRender. V.G.S. is an Investigator of the Howard Hughes Medical Institute.

## Author contributions

Conceptualization, P.L., G.A., and V.G.S.; Methodology, P.L., G.A., C.-J.G., S.A.A., S.R., and V.G.S.; Investigation P.L., C.-J.G., A.S., and T.Y.; Computational analyses, P.L., G.A., and C.W.; Resources, W.B., M.A., S.J., A.C., and M.P.; Funding acquisition and Supervision, V.G.S.; Writing - original draft, P.L., G.A., and V.G.S.; Writing - review & editing, P.L., G.A., and V.G.S. with input from all authors.

## Competing interest statement

S.A.A. has been a consultant and/or shareholder for Neomorph Inc., C4 Therapeutics, Accent Therapeutics, Stelexis Biosciences, AstraZeneca, Novartis, and Hyku Therapeutics. S.A.A. has received research support from Janssen and Syndax. SAA is an inventor on a patent application related to MENIN inhibition WO/2017/132398A1. V.G.S. is an advisor to Ensoma, Cellarity, and Beam Therapeutics, unrelated to this work. There are no other relevant declarations to report.

## Data availability

Raw and processed data have been deposited at GEO, and will be available at the time of publication. All source data used for figure generation with Graphpad Prism are included in Data S1. Individual-level genotypic and phenotypic data from the UK Biobank are available to qualified researchers upon application through the UK Biobank resource (www.ukbiobank.ac.uk/). Raw DMS-seq sequencing data generated in this study have been deposited in Zenodo (10.5281/zenodo.18903315). The GEO accession numbers of human HSC bulk-RNAseq and AML cell line ChIRP-RNAseq datasets are included in the key resources table and are accessible at GEO under accessions GSE301055 and GSE301056, respectively. No original code was generated in this study.

## Materials and Methods

### CD34^+^ HSPC culture

Enriched CD34^+^ HSPCs from G-CSF mobilized peripheral blood cells were purchased from Fred Hutch Cancer Institute. Cryopreserved HSPCs were rapidly thawed in a 37°C water bath, followed by serially dilution with thawing media (1× PBS supplied with 1% FBS). Thawed cells were cultured at 37°C in StemSpan Serum-Free Expansion Medium (SFEM) II supplemented with 1% L-Glutamine (200 mM), 1% Penicillin-Streptomycin (10,000 U/mL), 1× CC100 cytokine cocktail (containing FLT3L, stem cell factor (SCF), IL-3, and IL-6), 100 ng/mL thrombopoietin (TPO), and 35 nM UM171 (Stem Cell Technologies). Cord blood (CB) was acquired from Dana-Farber Cancer Institute’s Pasquarello Tissue Bank for CB CD34^+^ HSPCs selection. Briefly, CD34^+^ HSPCs were pre-enriched using Ro-setteSep™ Human Cord Blood CD34 Pre-Enrichment Cocktail (#15896C) and the sample was layered onto Ficoll-Paque for density gradient centrifuging. The pre-enriched fraction was then processed with EasySep™ Human CD34 Positive Selection Cocktail (#18096C) and purified CB CD34^+^ HSPCs were cryopreserved or used directly for downstream applications.

### Cell line culture

HEK293T cells were cultured in Dulbecco’s Modified Eagle’s Medium (DMEM) supplemented with 10% FBS and 1% penicillin/streptomycin. K562 cells were cultured in Iscove’s Modified Dulbecco’s Medium (IMDM) supplemented with 10% FBS, and 1% Penicillin-Streptomycin. MOLM13 cells were cultured in RPMI 1640 Medium supplemented with 20% FBS, and 1% Penicillin-Streptomycin^67^. OCI-AML4 cells were cultured in Minimum Essential Medium α (MEM α) supplemented with 20% FBS, and 1% Penicillin-Streptomycin^68^. MUTZ-3 cells were cultured in MEM α supplemented with 20% FBS, 20% conditioned medium of cell line 5637, and 1% Penicillin-Streptomycin^69^. All cell lines were cultured according to the supplier’s recommended protocols and maintained at appropriate confluency throughout the experiments.

### AML patient-derived cell *ex vivo* culture

We obtained cells from AML patients that we were able to maintain and give rise to patient-derived xenografts (PDXs) from which cells could be obtained and cultured. The CBAM-68552 PDX was initially obtained from the Public Repository of Xenografts^70^ and CBSK-17D was a kind gift from R. Levine, Memorial Sloan Kettering Cancer Center, to the Armstrong Laboratory. PDX specimens used in this study were previously expanded and banked in the Armstrong Laboratory. Cells derived from PDXs were thawed, followed by depletion of mouse cells using the Milltenyi Biotec Mouse Cell Depletion Kit. Thereafter, cells were cultured in the following media: StemSpan SFEM II, supplemented with 10 ng/mL Human Recombinant IL-3, 10 ng/mL Human Recombinant IL-6, 10 ng/mL Human Recombinant G-CSF, 25 ng/mL Human Recombinant TPO, 50 ng/mL Human Recombinant SCF, 50 ng/mL Human Recombinant Flt3/Flk-2 Ligand, 0.5 µM SR1, and 0.2% Primocin.

### Mouse xenotransplantation models

All the mouse procedures were conducted under the approval and ethical guidance of the Institutional Animal Care and Use Committee (IACUC) at Boston Children’s Hospital. CB CD34^+^ HSPCs were nucleofected with CRISPR/Cas9 and CBE RNPs separately, including *AAVS1* KO, *HOTSCRAMBL* KO, *AAVS1* CBE, and *HOTSCRAMBL*^rs17437411^. Two days post-nucleofection, a small fraction of cells was harvested to test the editing efficiency and assess the percentage of CD34^+^ cells in culture. A total of 200,000 CD34^+^ HSPCs were injected into each immunodeficient NOD.Cg-*Kit*^*W41J*^*Tyr*^+^*Prkdc*^*scid*^ *Il2rg*^*tm1Wjl*^/ThomJ (NBSGW) mouse (IMSR_JAX#026622) via tail vein. After transplantation, mice were supplied with autoclaved water containing sulfatrim antibiotics and refreshed weekly to prevent infections. To monitor the chimerism, mice were anesthetized by isoflurane, then peripheral blood was collected in EDTA-coated capillary tubes at week 4, 8, 12, and 16 via retro-orbital sinus bleeding, and antibiotic ophthalmic ointment was applied after the bleeding. Bone marrow (BM) and spleen were harvested 16 weeks after transplantation for analysis of engraftment, lineage chimerism, and *HOTSCRAMBL* editing efficiency post-transplantation, following a previously reported protocol with modifications^71^. In brief, mice were euthanized by CO_2_, and total BM cells were obtained from ilium, femurs, and tibias. Spleen cells were isolated by mechanical dissociation. Before red blood cells (RBC) lysis, 1 mL of BM cells was taken for erythroid engraftment analysis. RBCs were depleted using RBC lysis buffer for both BM and spleen samples. The remaining cells were resuspended in FACS buffer (1× PBS supplemented with 2% FBS and 2 mM EDTA) and filtered through a 70 µm cell strainer to obtain single-cell suspensions of RBC-free BM and spleen cells for further analyses.

### Genetic associations at the *HOXA* locus

Genome-wide significant common variants associated with blood cell traits at the *HOXA* cluster were identified from a meta-analysis of blood cell traits^23^ to highlight lead single nucleotide polymorphisms (SNPs) at independent haplotypes. For rs35355140, phenome-wide associations were based on a large multi-phenotype GWAS in UK Biobank (UKB) examining 2,419 traits^24^. Associations with clonal phenotypes were examined using GWAS summary statistics for clonal hematopoiesis of indeterminate potential (CHIP) subtypes^25^. For telomere length, the phenotype used was leukocyte telomere length measured in UKB participants^26^ (Data Field 22192). As telomere length varies across ancestries, we restricted this analysis to European-ancestry individuals in the UKB, and quality control (QC) and normalization were performed as previously described^27^. Association with rs35355140 was assessed using linear regression adjusted for age, sex, and the first 10 genetic principal components.

For association with myeloproliferative neoplasms (MPNs), a meta-analysis was performed across UKB, Finngen, Vanderbilt BioVU, 23andMe, CIMBTR, and Wellcome Trust cohorts. The MPN phenotype was curated based on hospital codes and self-reported diagnoses for polycythemia vera, essential thrombocythemia, chronic myeloid leukemia, and myelofibrosis, as previously described^28^. Individual cohort-level GWAS were performed, adjusting for age, sex and top 5-10 genetic principal components as covariates into a logistic regression model. To estimate MPN risk by rs35355140 genotype, a fixed-effect size inverse variance-weighted meta-analysis was performed across these cohorts, additionally including association for detectable *JAK2*-mutant CHIP in UKB.

### Fine-mapping

Fine-mapping was performed at the sentinel SNP rs35355140, using blood cell trait associations available from the GWAS catalogue for monocyte count [GCST90002393], neutrophil count [GCST90002398] and platelet count [GCST90002402], and linkage disequilibrium (LD) matrices for UKB participants of British ancestry^76^. Variants with minor allele frequency (MAF) > 0.001 within 500 kB of the sentinel SNP rs35355140 were fine-mapped using SuSiE^77^ CARMA^78^, allowing up to 10 causal variants and reporting 95% credible sets for each method. Both methods retrieved the same credible set consisting of 4 SNPs (detailed in **Table S2**). Candidate SNPs at this locus were functionally annotated with their genomic context predicted splicing effects (using SpliceAI^59^).

### Cytosine base editing (CBE) protein purification

A modified version of the previously described base editor protein purification protocol was followed^79^ using the bacterial codon optimized TadCBE expression vector^80^. Briefly, the TadCBE vector was transformed into BL21 Star DE3 competent cells (Thermo Fisher C601003). The bacteria were grown in Terrific Broth media until they reached an optical density at 600 nm between 1.5 and 1.8. The culture was cooled on ice for 1 hour (hr) before protein expression was induced with the addition of a solution of 30% L-rhamnose. The culture was then incubated at 18°C for 24 hrs. After incubation, the bacteria were pelleted and flash frozen for storage at -80°C. On the day of protein purification, the bacteria pellet was thawed and vortexed with 60 mL of lysis buffer (20 mM HEPES buffer, 2 M NaCl, 1 mM TCEP, 10% glycerol, Roche Protease Inhibitor, DNase I) for every 0.5 L of bacteria thawed. The bacteria were sonicated for 18 minutes (min) using a 1/4” (6 mm) probe and then centrifuged at 20,000×g for 30 min. The lysate was crudely purified on a 5 mL Cytiva HisTrap FF column using a syringe pump system and following the standard Cytiva manual protocol for this column. The column was washed with a wash buffer (20 mM HEPES buffer, 2 M NaCl, 25 mM imidazole, 1 mM TCEP, 10% glycerol) and the protein was eluted with an elution buffer (20 mM HEPES buffer, 500 mM imidazole, 1 mM TCEP, 10% glycerol). The crude protein was concentrated to half of the original elution volume using a Amicon 100 kDa MWCO centrifugal filter column (MilliPore Sigma UFC9100). The crude protein was diluted 2× in a low salt buffer (20 mM HEPES buffer, 1 mM TCEP, 10% glycerol) and added to a 5 mL Cytiva HiTrap SP HP column. The column was run using a syringe pump system and following the standard Cytiva manual run protocol for this column. A gradient of low salt buffer to high salt buffer (20 mM HEPES buffer, 2 M NaCl, 1 mM TCEP, 10% glycerol) was applied to the column to elute the protein and ten fractions were collected. The fractions were run with SDS-PAGE to identify those containing the desired protein. The fractions with TadCBE were combined and concentrated using Amicon 100 kDa MWCO centrifugal filter columns until a protein concentration of ∼60 µM was reached. The concentrated protein was aliquoted, and flash frozen for storage at -80°C.

### Cytosine base editing and CRISPR/Cas9 editing

Nucleofection was performed using the Lonza 4D-Nucleofector system with the P3 Primary Cell 4D-Nucleofector (20 µL cuvette) for CD34^+^ HSPCs and primary AML patient-derived cells on day 2 after thawing, and the SF Cell Line 4D-Nucleofector (100 µL cuvette) for leukemia cell lines (including MOLM13, OCI-AML4, MUTZ-3, and K562) at approximately 75% confluence. Before nucleofection, cells were harvested and washed three times with DPBS. For CRISPR/Cas9 editing, sgRNAs were designed using the Synthego knockout guide design tool (https://design.synthego.com/#/) (**Table S10**). Ribonucleoprotein (RNP) complexes were assembled by mixing 100 pmol of sgRNA with 100 pmol of Cas9 protein in 20 µL of Lonza nucleofection buffer, followed by incubation at room temperature for 10-20 min. Cells were then resuspended in a nucleofection buffer supplemented with IDT nucleofection enhancer at a 20:1 ratio and mixed with the assembled RNP complexes (scaled up for cell line editing). Cytosine base editing (CBE) was performed as previously described^79^, with sgRNA validated using the BE-Hive tool (https://www.crisprbehive.design/)^81^ (**Table S10**). In brief, CBE RNP complexes were assembled by mixing 157 pmol of sgRNA with 20 µg of purified CBE-Cas9NG protein in 20 µL of Lonza nucleofection buffer (scaled up for cell line editing), and incubated at room temperature for 20 min. Cells were then resuspended in a nucleofection buffer and combined with the assembled RNP complexes.

For both CRISPR/Cas9 and CBE editing, program DZ-100 was used for human CD34^+^ HSPCs and primary AML patient-derived cells, and DS-150 for leukemia cell lines. After 72 or 96 hrs post-nucleofection, cells were harvested, and genomic DNA was extracted using the DNA Isolation Micro Kit. DNA fragments flanking the editing sites were PCR-amplified with primers listed in Table S11. For dual sgRNA-mediated deletions, editing efficiency was assessed by agarose gel electrophoresis. For single guide knockouts or CBE, editing outcomes were evaluated by Sanger sequencing or next-generation sequencing (NGS). CRISPR/Cas9 editing efficiency and frameshift composition were analyzed using the Inference of CRISPR Edits (ICE) tool (https://ice.synthego.com/#/). Base editing efficiency was quantified using EditR (https://moriaritylab.shinyapps.io/editr_v10/)^82,83^.

### Flow cytometric analysis

Flow cytometric analyses were performed based on standard protocols. Cells were washed three times with DPBS and resuspended in 100 µL FACS buffer (DPBS + 2% FBS + 2 mM EDTA) containing fluorescent dye or antibodies at desired dilution. The details of antibody color panels and usage were listed in **Table S9**. Stained cells were washed and resuspended in the FACS buffer for flow cytometric analysis. All data were acquired using BD LSRFortessa (DB Biosciences) and analyzed with FlowJo software.

For human HSC profiling, cultured CD34^+^ HSPCs were stained with combinations of antibodies including CD34, CD45RA, CD90, CD133, EPCR and ITGA3. In leukemia cell differentiation assay, the myeloid differentiation markers CD11b and CD14 were used for evaluating monocytic and granulocytic differentiation in primary AML patient-derived cells (CBSK-17D and CBAM-68552) and AML cell lines (MOLM13 and OCI-AML4); CD34 and CD33 were used to assess progenitor status and early myeloid commitment in MUTZ-3; erythroid markers CD71 and CD235a were used to monitor erythroid differentiation in K562. In mouse xenotransplantation experiments, anti-human and anti-mouse CD45 antibodies were used to distinguish human chimerism in PB samples, and representative blood lineage markers were stained to analyze lineage outcomes in BM and Spleen (**Table S9**). Cell surface marker staining was carried out for 20-40 min at room temperature protected from light, followed by two DPBS washes.

Cell cycle analysis was performed using Vybrant™ DyeCycle™ Violet according to the manufacturer’s instructions. Briefly, cells were suspended in complete culture medium at a concentration of 0.5-1×10^6^ cells/mL. Dyecycle staining dye was added to a final concentration of 5 µM, followed by incubation at 37°C for 30 min in the dark. Cells were then analyzed directly without further wash.

For cell division assays, cells were labeled with CellTrace™ CFSE after nucleofection according to the manufacturer’s instructions. Cells were washed twice, then resuspended in DPBS containing 2 µM CFSE and incubated at 37°C for 10 min. Labeling was quenched by adding five volumes of DPBS containing 2% FBS. After washing, cells were resuspended in HSC medium and cultured under appropriate conditions. CFSE dilution was assessed over time to monitor cell division. Cell apoptosis was assessed by Annexin V and Propidium Iodide (PI) co-staining. Cells were washed and resuspended in 100 µL of Annexin V binding buffer containing 5 µL of Annexin V and 1 µg/mL of PI, followed by incubation for 20 min at room temperature in the dark.

### RNA fluorescence in situ hybridization (FISH) and Immunofluorescence (IF)

DIG- or Biotin-labeled antisense RNA probes were synthesized using the DIG or Biotin RNA Labeling Mix and T7 RNA Polymerase according to the manufacturer’s protocol. Briefly, DNA templates containing a T7 promoter were generated by PCR amplification using gene-specific primers and purified with Monarch^®^ PCR purification kit. *In vitro* transcription (IVT) was performed using T7 RNA polymerase at 37°C for 2 hrs in the presence of DIG or Biotin RNA Labeling Mix (Roche), followed by DNase I treatment to remove the DNA template. RNA was then purified by ethanol precipitation supplemented with 1/10 volume of 3 M sodium acetate. Probe concentrations were determined by NanoDrop and stored at -80°C.

RNA FISH was performed as previously described^84,85^ with minor modifications. Human CD34^+^ or sorted CD34^+^CD45RA^−^CD90^+^ HSPCs were cytospun (Thermo Scientific Cytospin 4 Centrifuge) on glass slides followed by a fixation in 4% PFA with 10% acetic acid for 15 min at room temperature, permeabilized in 0.5% Triton X-100 with 2 mM Ribonucleoside Vanadyl Complex (RVC), and dehydrated in 75% ethanol. Cells were hybridized overnight at 50°C with denatured DIG-labeled *HOTSCRAMBL* probes and Biotin-labeled *HOXA9* pre-mRNA probes (sense probe used as negative control) in 50% formamide in 2×SSC. After three times washing in Sal I buffer (50% formamide, 0.1×SSC, 0.1% SDS), cells were incubated at room temperature with sheep anti-DIG antibody (Roche, 1:400 in blocking buffer) and/or mouse anti-Biotin antibody (Roche, 1:400 in blocking buffer) under parafilm in a humidified chamber. Following three washes with Sal II buffer (2×SSC containing 8 % formamide), Alexa Fluor 647-conjugated donkey anti-sheep secondary antibody (1:800) and/or Alexa Fluor 594-conjugated donkey anti-mouse secondary antibody, was applied for 1 hr in the dark.

IF was performed followed by FISH staining, in brief, cells were washed three times with 1× PBS at room temperature. And cells were incubated with a blocking buffer (1×PBS supplemented with 0.5% Triton X-100, 3% BSA, and 5% donkey or goat serum) for 1 hr at room temperature. Then, primary antibodies were diluted in the blocking buffer (SRSF2, 1:1000; hnRNPA1, 1:800) and applied to the cells overnight at 4°C. After primary incubation, cells were washed three times with 1×PBS and incubated with the Alexa Fluor conjugated secondary antibodies goat anti-mouse (594nm, hnRNPA1) or goat anti-rabbit (488nm, SRSF2), at 1:1000 dilution in 1% BSA/PBS for 2 hrs at room temperature in the dark. Nuclei were counterstained with DAPI, and slides were mounted before imaging. Fluorescence was acquired with a Zeiss LSM 980 Airyscan2 confocal microscope.

### Real time quantitative PCR (qPCR) and Reverse transcription PCR (RT-PCR)

Total RNA from sorted or unsorted cells was isolated using the Total RNA Purification Micro Kit (Norgen Biotek) with DNase I treatment according to the manufacturer’s instructions. For cDNA synthesis, an equal amount of RNA was reverse transcribed using the iScript cDNA Synthesis Kit (Bio-Rad).

The synthesized cDNA was diluted 1:10, and 2 µL of diluted cDNA was used as input for qPCR using iQ SYBR Green Supermix (Bio-Rad) on CFX96 or CFX384 systems (Bio-Rad). Relative expression levels were normalized to housekeeping genes and calculated using the ΔΔCt method, with control samples serving as the reference.

PCR amplification of cDNA was performed using gene-specific primers in a thermal cycler under standard cycling conditions. Multiple PCR cycle numbers were tested for each gene and primer pair, and cycles in the range of exponential amplification phase were used for semi-quantitative analysis. PCR products were analyzed on a 1.2% agarose gel containing SYBR Safe DNA Gel Stain and visualized using a ChemiDoc imaging system (Bio-Rad).

The measurement of *HOTSCRAMBL* and *HOXA9* pre-mRNA copy number was adapted from a previously described protocol^86^ with minor modifications. Briefly, Linearized pGEM-T-*HOTSCRAMBL* or pGEM-T-*HOXA9* plasmids were serially diluted and used to establish standard curves for absolute quantification. Plasmid copy numbers corresponding to each dilution were estimated using the DNA/RNA Copy Number Calculator (http://endmemo.com/bio/dnacopynum.php). For endogenous measurements, total RNA was isolated from 1×10^6^ MOLM13 cells and reverse-transcribed into cDNA. Quantitative PCR was then performed, and absolute copy numbers of *HOTSCRAMBL* and *HOXA9* pre-mRNA were interpolated from the respective standard curves.

### Multiplexed Error-Robust Fluorescence in situ Hybridization (MERFISH)

MERFISH experiments were performed based on previously described protocols with minor modifications^87,88^. The probe pool targeting *HOTSCRAMBL* was designed using the MERFISH probe design pipeline (https://github.com/xingjiepan/MERFISH_probe_design) and synthesized by Integrated DNA Technologies (IDT) as oPools™ Oligo Pools (**Table S10**). Each encoding probe contained a ∼30-nucleotide sequence complementary to the target RNA flanked by 5′ and 3′ adapter overhangs. The probe pools used in this study included smFISH_*HOTSCRAMBL*_[Stv_22, Stv_23]. Adapter probes complementary to the encoding probe overhangs were synthesized by IDT, including Stv_22_2xStv79rc (TTCCGCGTCATGTACCGGTTTGCGAACTGTCCGGCTTTCATGCGAACTGTCCGGCTTTCA) and Stv_23_2xStv1rc (GTGGGCTGCTGCGATTTCACGATCCGATTGGAACCGTCCCGATCCGATTGGAACCGTCCC). Fluorescent readout probes labeled with Alexa Fluor 647 (Stv1) (/5Alex647N//iThioMC6-D/GGGACGGTTCCAATCGGATC) and ATTO565 (Stv79) (/5ATTO565N//iThioMC6-D/TGAAAGCCGGACAGTTCGCA) were used for signal detection. These probes enabled sequential hybridization of adapter and fluorescent readout probes to visualize individual RNA molecules.

Human CD34^+^ HSPCs subjected to gene editing (control, AAVS1 knockout, or HOTSCRAMBL knockout) were deposited onto MAS-GP adhesive glass slides using Cytospin centrifugation. Cells were fixed with 4% paraformaldehyde for 10 min at room temperature, washed with RNase-free PBS, and permeabilized with 0.5% Triton X-100 supplemented with RNase inhibitor. The *HOTSCRAMBL* probe pool was hybridized in 2×SSC containing 30% formamide, 10% dextran sulfate, and 0.1% yeast tRNA with RNase inhibitor. Samples were incubated at 37°C for 24 hrs, followed by washes in 30% formamide/2×SSC to remove excess probes. Adapter probes (100 nM) were hybridized in 30% formamide/2×SSC for 15 min at room temperature, followed by incubation with fluorescent readout probes (20-25 nM) labeled with Alexa Fluor 647 or ATTO565. Cells were counterstained with DAPI and mounted in an antifade imaging buffer. Fluorescence images were acquired using a confocal fluorescence microscope, and individual RNA molecules were detected as discrete fluorescent puncta.

### Subcellular fractionation

Cytoplasmic and nuclear RNA from CD34^+^ HSPCs was isolated as previously described^86^. Briefly, 5×10^5^ cells were washed with ice-cold DPBS and lysed in 100 µL of lysis buffer, containing 10 mM Tris-HCl (pH 8.0), 140 mM NaCl, 1.5 mM MgCl_2_, 0.5% Igepal, 2 mM RVC on ice for 5 min. One-fifth of the lysate was collected for total RNA extraction with Trizol reagent. The remaining lysate was centrifuged at 1,000×g for 3 min at 4°C to collect nuclei (pellet) and cytoplasm (supernatant). The cytoplasmic fraction was further clarified by centrifugation at 16,000×g for 10 min at 4°C before RNA extraction. The nuclear pellet was washed once with lysis buffer and once with lysis buffer containing 0.5% deoxycholic acid, then resuspended in 100 µL lysis buffer and extracted with Trizol reagent. An equal amount of RNA was used for cDNA synthesis and qPCR.

### Colony Forming Unit (CFU) assay

CD34^+^ HSPCs were plated after three days post-nucleofection at a density of 500 cells/mL methylcellulose medium (H4034), and primary AML patient-derived cells were plated four days post-nucleofection at a density of 5,000 cells/mL methylcellulose medium (H4034) following the manufacturer’s instructions. Approximately 1 mL of the cell with culture media was dispensed into each well of a SmartDish (StemCell Technologies) using a blunt-end needle and syringe. Plates were gently rocked to evenly distribute the medium and placed in humidified trays. Cultures were incubated at 37°C and colonies were counted on day 14. For serial replating CFU assays, 5,000 CD34^+^ HSPCs from long-term xenotransplant mouse recipients were plated in methylcellulose medium (H4434). After 10 days of primary plating, the colonies were counted, and then harvested for secondary replating with an equal number of cells (based on primary colony yield). All the colonies were imaged by using StemVision system (StemCell Technologies) and colony details were determined using STEMvision™ Analyzer (StemCell Technologies) and STEMvision™ Colony Marker Applications (StemCell Technologies).

### LNA GapmeR knockdown

Antisense LNA GapmeRs (Qiagen) were used to knock down target RNAs in primary human CD34^+^ HSPCs^89,90^. GapmeRs were designed using Qiagen’s online tool (https://geneglobe.qiagen.com/us/customize/designers/antisense-lna-gapmers/configure) and diluted in TE buffer to a 1 µM working concentration. Cells (5×10^5^) were electroporated using the 4D-Nucleofector^®^ X Unit (Lonza) and the P3 Primary Cell Kit (V4XP-3032), following the manufacturer’s instructions. Briefly, cells were resuspended in a nucleofector solution containing 75 nM GapmeR and electroporated using program EO-100. Immediately after electroporation, cells were transferred into pre-warmed StemSpan II medium and incubated at 37°C with 5% CO_2_. Total RNA was harvested 48-72 hrs post-transfection for assessment of knockdown efficiency.

### SnoVector overexpression

To overexpress nuclear-localized RNAs, the snoVector was utilized as described previously^29^. Briefly, the *HOTSCRAMBL* and *HOTSCRAMBL*^rs17437411^ full length RNA was cloned into the snoVector backbone, enabling stable nuclear expression. For delivery into primary human CD34^+^ HSPCs, the snoVector plasmid was electroporated using the 4D-Nucleofector™ X Unit (Lonza) and the P3 Primary Cell Kit (V4XP-3032), following the manufacturer’s recommendations. Typically, 0.5 ×10^6^ CD34^+^ HSPCs were resuspended in 20 µL of supplemented P3 buffer containing 2-3 µg plasmid DNA, and electroporated using program EO-100. After electroporation, cells were immediately transferred into pre-warmed StemSpan II medium supplemented with appropriate cytokines and incubated at 37°C with 5% CO_2_. RNA expression was evaluated 48-72 hr post-transfection by qPCR.

### Bulk RNA-seq analysis

After three days CD34^+^ HSPC editing, CD34^+^CD45RA^-^CD90^+^ HSCs were sorted out to extract the RNA. The RNA sequencing library was prepared using the SMART-Seq® mRNA LP Kit (Takara) according to the manufacturer’s instructions. Briefly, 30 ng of total RNA from each condition was used for cDNA synthesis and amplification following the SMART-Seq protocol. Quality and concentration of cDNA libraries were assessed by using an Agilent 2100 Bioanalyzer. Libraries were sequenced on the Illumina NextSeq™ 2000 platform using the P4 XLEAP-SBS™ 300-cycle kit.

Raw sequencing FASTQ files were first assessed for quality using FastQC (v0.12.1). Adapter sequences and low-quality bases were removed with Trimmomatic (v0.39)^91^. Cleaned reads were aligned to the human reference genome (GRCh38) using STAR aligner (v2.7.11b)^92^ with default parameters. Transcript assembly and quantification were performed using StringTie (v3.0.0)^93^. To investigate splicing dynamics, splice junction usage was analyzed using SplAdder (v3.0.5)^66,94^. Gene-level expression counts were calculated using featureCounts (v2.0.6)^95^ from the Subread package. Differential gene expression analysis was conducted in R (4.4) using the DESeq2 package (v1.48.1)^96^ (**Table S3**). Data visualization, including volcano plots and heatmaps, was performed using ggplot2 (v3.5.1) in R platform. Genomic track visualization was carried out using the Integrative Genomics Viewer (IGV) (v2.18.4)^97^.

### Western blotting

Whole cell lysates were prepared from primary CD34^+^ HSPCs, flow-sorted CD34^+^CD45RA^−^CD90^+^ subsets, and leukemia cell lines. Cells were lysed in ice-cold RIPA buffer (Thermo Fisher) supplemented with protease inhibitor cocktail (Roche) for 30 min on ice. Lysates were centrifuged at 14,000×g for 15 min at 4°C to remove debris. Protein concentrations were determined using the Pierce™ BCA Protein Assay Kit (Thermo Fisher) according to the manufacturer’s instructions.

Equal amounts of protein (20-40 µg per sample) were mixed with 4× Laemmli loading buffer (Bio-Rad) containing 10% β-mercaptoethanol, and boiled at 98°C for 10 min to denature proteins. Samples were loaded on 4-15% Mini-PROTEAN® TGX™ Precast Gels (Bio-Rad) and transferred onto PVDF membranes. Membranes were blocked with 5% BSA in TBST (Tris-buffered saline with 0.1% Tween-20) for 1 hr at room temperature, followed by incubation with primary antibodies overnight at 4°C. After washing, membranes were incubated with HRP-conjugated secondary antibodies and developed using ECL substrate (Bio-Rad). Signal intensity was quantified using ImageJ software and normalized to internal loading controls.

### Lentivirus packaging and transduction

Human *HOXA9* coding sequence was cloned into pLVX-EF1a-ZsGreen vectors by using In-Fusion^®^ Snap Assembly kit, and the primers were designed by using their online tool (https://www.takarabio.com/learning-centers/cloning/primer-design-and-other-tools). Lentiviral particles were produced in HEK293T cells using a standard three-plasmid system (pLVX transfer vector, psPAX2 packaging plasmid, and pMD2.G envelope plasmid) with a 2:1.5:1 molar ratio. Transfection was performed using Lipofectamine 3000 (Invitrogen). Viral supernatants were collected 48 hrs post-transfection, filtered through 0.45 µm filters, and concentrated with ultra-high speed centrifuge. Viral titers were estimated by using Lenti-X™ Provirus Quantitation Kit (Takara) in K562 cells as per manufacturer’s instructions.

For transduction of human HSPCs, cells were resuspended in serum-free StemSpan II medium with 8 µg/mL cyclosporin H, incubated with virus at equal MOI 20-50, and spinfected at 450×g for 90 min at room temperature. Cells were then cultured at 37°C for >48 hrs before downstream analysis.

### Chromatin Isolation by RNA Purification (ChIRP)

ChIRP was performed to isolate lncRNA-associated DNA/RNA complexes following a modified protocol^98^. Biotinylated antisense DNA probes (20 nt) complementary to the target lncRNA were designed using the Biosearch Technologies online tool, ChIRP Probe Designer (https://www.biosearchtech.com/chirp-designer) followed the instruction. Probes were synthesized with a 3’ Biotin-TEG modification (Biosearch Technologies).

For each sample, 1×10^7^ MOLM13 or OCI-AML4 cells were fixed in 10 mL of freshly prepared 1% glutaraldehyde at room temperature for 10 min, then quenched with 125 mM glycine for 5 min. Fixed cells were washed in ice-cold DPBS, flash-frozen, and stored at -80°C. Fixed cell pellets were thawed and resuspended in lysis buffer (1 mL per 10^7^ cells) supplemented with 1×PMSF (1 mM), 1× protease inhibitor (Add 1 tablet into 1 mL ddH_2_O to make 50× stock), and RNase inhibitor (0.2 U/µL), then sonicated using a Covaris E220 ultrasonicator with optimized program (PIP: 150W, DF: 15%, CPB: 400, Time: 30-60 min) to obtain 100-500 bp chromatin fragments. The efficiency of fragmentation was verified by agarose gel electrophoresis after proteinase K digestion and DNA purification (Qiagen). For each ChIRP reaction, lysates were incubated with biotinylated probes (100 pmol per 1 mL lysate) in Hybridization Buffer at 37°C for 4 hrs with rotation. Pre-washed Dynabeads™ MyOne™ Streptavidin C1 beads (100 µL per 100 pmol probe) were added and incubated for an additional 30 min. Beads were washed five times at 37°C using Wash Buffer to reduce nonspecific binding. For RNA isolation, beads were incubated in RNA PK Buffer with proteinase K at 50°C for 45 min, followed by heat revere cross-linking at 95°C for 10 min. RNA was extracted using TRIzol reagent and purified with the miRNeasy Mini Kit (Qiagen). Residual DNA was removed using the TURBO DNA-free™ Kit (AM1907, ThermoFisher Scientific). Enriched RNA was quantified for qPCR or RNA sequencing.

For ChIRP-RNA seq, the RNA integrity was assessed with a Bioanalyzer (Agilent) and quantified with Qubit 4 Fluorometer (Thermo Scientific), and libraries were constructed using the SMART-Seq® Total RNA Pico Input Kit (Takara) according to the manufacturer’s instructions. Indexed libraries were pooled and sequenced on the Illumina NextSeq 1000 platform using the NextSeq™ 1000/2000 P2 XLEAP-SBS™ Reagent Kit (100 cycles, Illumina). For intron read analysis, aligned BAM files were generated using STAR, and transcript annotations (GTF) were obtained using StringTie2. *HOXA9* intron coverage was subsequently quantified using BEDtools (v2.31.1)^99^.

### Comprehensive identification of RNA-binding proteins by mass spectrometry (ChIRP-ms)

ChIRP-MS was performed as previously described with modifications^36^. Briefly, 1×10^7^ cells were crosslinked using 10 mL of 1% formaldehyde (Thermo Scientific) at room temperature for 30 min and quenched with 125 mM glycine. Fixed cell pellets were thawed and resuspended in lysis buffer (1 mL per 10^7^ cells) supplemented with 1× PMSF, 1× protease inhibitor, and RNase inhibitor, then sonicated using a Covaris E220 ultrasonicator with optimized program (PIP: 150W, DF: 10%, CPB: 200, Time: 30-60 min) to obtain 200-700 bp chromatin fragments. Biotinylated antisense DNA probes targeting the lncRNA of interest were designed using Biosearch Technologies’ online tool and hybridized to lysates at 37°C for 12-16 hrs. Probe-bound complexes were captured using Dynabeads™ MyOne™ Streptavidin C1 (Thermo Fisher), followed by stringent washing steps. Bound proteins were eluted using a biotin elution buffer and precipitated with trichloroacetic acid (TCA). Eluted proteins were solubilized in Laemmli buffer, boiled, and resolved by SDS-PAGE using NuPAGE Bis-Tris gels. Protein bands were visualized using Pierce™ Silver Stain for Mass Spectrometry (Thermo Scientific) according to the manufacturer’s protocol. Protein identification was carried out at Thermo Fisher Scientific Center for Multiplexed Proteomics core.

### Dimethyl sulfate mutational profiling with sequencing (DMS-MaPseq)

DMS-MaPseq was performed on two replicates of each *HOTSCRAMBL* variant. Additionally, no DMS control (excluding DMS below) was performed on each variant. In brief, DNA templates of *HOTSCRAMBL* and *HOTSCRAMBL*^rs17437411^ were amplified by CloneAmp HiFi PCR Premix by adding a 5-prime T7 promoter on the forward primer and cleaned using the Monarch Spin PCR & DNA Cleanup Kit. RNA was generated using HiScribe T7 High Yield RNA Synthesis Kit incubated at 37°C for 2 hrs. The DNA template was removed using TURBO DNAse (ThermoFisher Scientific AM2238) at 37°C for 1 hr. RNA was purified using the Monarch Spin RNA Cleanup Kit.

For *in vitro* RNA folding and DMS-MaPseq, refolding buffer (0.4M Sodium cacodylate), 6 mM MgCl_2_ was preheated to 37°C. Purified *in vitro*-transcribed RNA was suspended to 0.5 µg/µL in 10 µL of H_2_O, incubated at 95°C for 1 min, immediately mixed with 87 µL of preheated refolding buffer, and incubated at 37°C for 30 min. 3% (3 µL) DMS was added to each reaction and incubated at 37°C for 5 min before being quenched with 60 µL β-Mercapethanol. RNA was isolated using Monarch Spin RNA Cleanup Kit, and each sample was reverse transcribed using Induro Reverse Transcriptase according to manufacturer protocol using primers in **Table S11**. cDNA was amplified using CloneAmp HiFi PCR Premix for 25 cycles. Amplicons were purified using Monarch Spin PCR & DNA Cleanup Kit and sequencing library was prepared using NEBNext UltraExpress DNA Library Prep Kit and NEBNext Multiplex Oligos. Samples were pooled and sequenced on an Illumina NextSeq 1000 platform (2× 150 bp paired-end reads) according to the manufacturer protocol.

DMS-MaPseq sequencing data was processed using SEISMIC v24.3 (https://github.com/rouskinlab/seismic-rna) to map DMS reactivity, cluster alternative structures, and evaluate secondary structure models^100^. Prior to clustering and structure mapping replicates were pooled for analysis. Structure model visualizations were generated using VARNA^101^. Nucleotide 403 was masked for clustering and structure prediction due to disproportionate DMS reactivity. DMS signals were normalized using Yeo-Johnson power transformation for direct comparison.

### RNA Immunoprecipitation (RIP)

RNA immunoprecipitation was performed as previously described^86,102^, with some modifications. Briefly, cells were washed with cold PBS and lysed in RIP buffer (50 mM Tris-HCl pH 7.4, 150 mM NaCl, 0.5% Igepal, 1 mM PMSF, protease inhibitors, and 2 mM RVC), followed by three cycles of gentle sonication (Covaris). Lysates were clarified by centrifugation and precleared with Dynabeads Protein G (Invitrogen). Supernatants were incubated with 5 µg SRSF2 antibody (Proteintech) or Rabbit IgG control for 4 hrs at 4°C. Immune complexes were washed with high-salt buffer and RIP buffer, and RNA was extracted using TRIzol reagent. Enrichment of *HOTSCRAMBL* and *HOXA9* was measured by RT-PCR or qPCR (normalized to input).

### Minigene splicing assay

The splicing of *HOXA9* was analyzed by minigene construct using pSpliceExpress vector^56^ as previously described^103^. In brief, genomic regions of *HOXA9* spanning exon 1 to exon 2 were PCR-amplified from human genomic DNA and cloned into the pSpliceExpress vector using seamless cloning with In-Fusion^®^ Snap Assembly. The recombinant plasmid was transformed into *stbl3* competent cells (Invitrogen) according to the manufacturer’s instructions. Bacterial colonies were screened by colony PCR, and positive clones were verified by Sanger sequencing. For transfection, 2-5 µg of plasmid DNA was introduced into MOLM13 cells using the SF Cell Line 4D-Nucleofector (Lonza) with the EO-100 program. Total RNA was extracted 2 days post-transfection using a Norgen RNA purification micro kit and reverse transcribed using the iScript cDNA Synthesis Kit (Bio-Rad). Splicing efficiency was evaluated by RT-PCR using exon-spanning primers. PCR products were resolved on 1.5% TAE agarose gels and visualized with SYBR™ Safe DNA Gel Stain.

### Quantification and Statistical Analysis

All experiments were performed with at least three independent biological replicates. For studies using primary human CD34^+^ HSPCs, biological replicates were defined as samples derived from distinct donors. For xenotransplantation experiments, each biological replicate corresponds to an individual recipient mouse, with a minimum of five animals per group.

Data are presented as mean ± standard deviation (SD) unless otherwise noted. For comparisons between two groups, statistical significance was determined using a two-tailed unpaired Student’s t test. If the data were not normally distributed or variance homogeneity could not be assumed, the nonparametric Mann-Whitney U test was used as an alternative. For comparisons among three or more groups, Levene’s test was first applied to assess homogeneity of variances. If equal variances were assumed, one-way or two-way analysis of variance (ANOVA) was performed, followed by post hoc multiple comparison testing using Dunnett’s test (when comparing each group to a control group) or Tukey’s test (when comparing all groups pairwise). If Levene’s test indicated significantly different variances across groups, a nonparametric Kruskal-Wallis test was employed, followed by the Dunn’s test for post hoc pairwise comparisons.

All statistical analyses were conducted using GraphPad Prism (V10) or R (v4.4.0 or later). Statistical significance was defined as \**P* < 0.05, \*\**P* < 0.01, \*\*\**P* < 0.001. Exact *P* values and the statistical tests used are indicated in the corresponding figure legends.

**Supplementary Figure 1:**
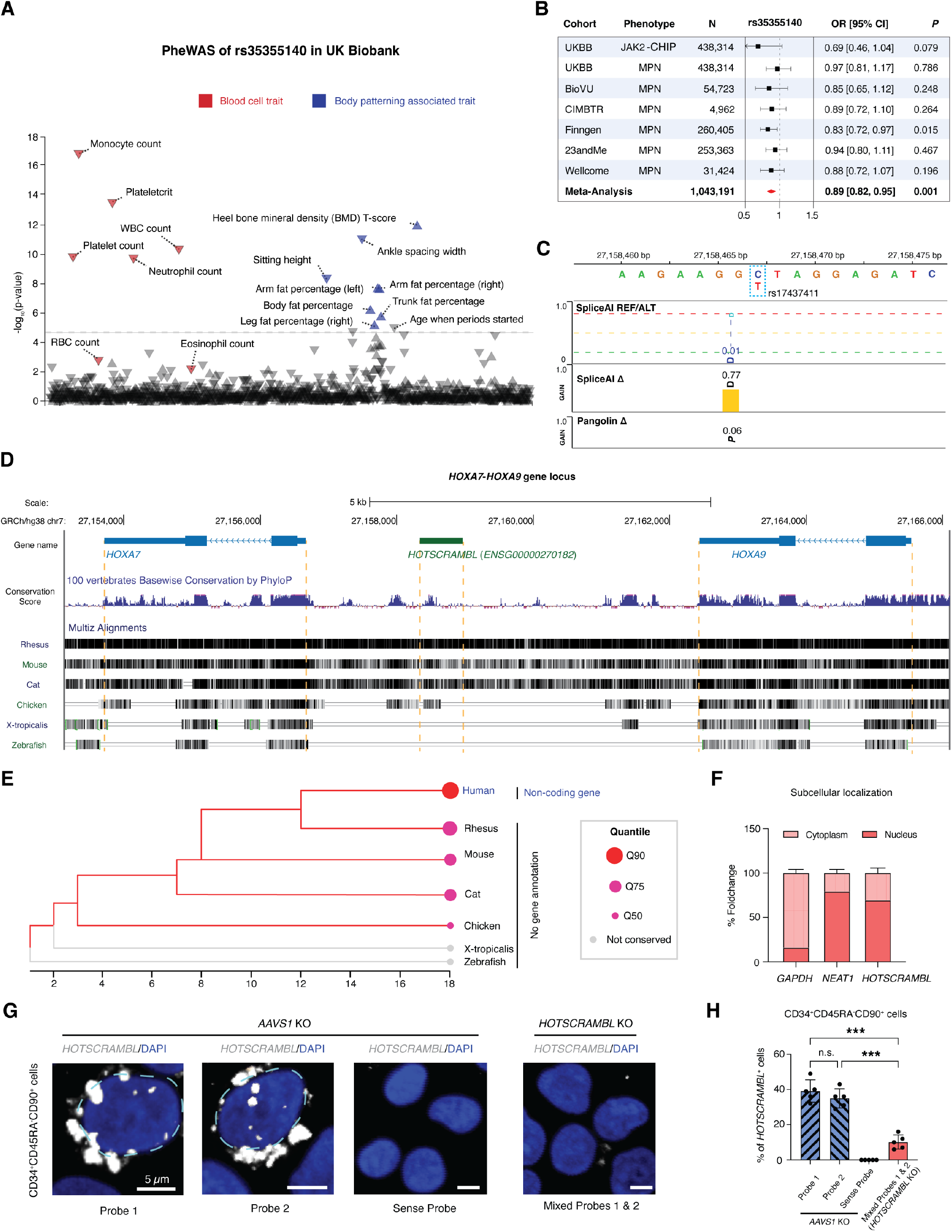
The human-specific, nucleus-enriched lncRNA *HOTSCRAMBL* harbors rs17437411, a common variant associated with multiple blood cell traits. **(A)** Phenome-wide association study (PheWAS) plot showing -log_10_(P) value associations for the rs17437411 variant in UK Biobank58, highlighting blood cell counts (red) and body patterning traits (blue). **(B)** Association of rs35355140 with myeloproliferative neoplasms (MPN) and JAK2-mutant clonal hematopoiesis phenotypes, across multiple cohorts and by meta-analysis. **(C)** Prediction of the splicing impact of rs17437411 based on SpliceAI59 and Pangolin Δ score60 models. **(D)** UCSC genome browser view of PhyloP conservation scores61 and multiple alignment across species at the *HOXA7* to *HOXA9* locus, gene body regions of *HOXA7, HOTSCRAMBL* (*ENSG00000270182*), and *HOXA9* are denoted in dashed orange lines. **(E)** Phylogenetic tree and conservation quantile ranks of *HOTSCRAMBL* region across representative vertebrates62. Bubble size represents the level of sequence conservation; species without similar sequences are shown in grey. **(F)** Percentage fold change of nuclear and cytoplasmic enrichment for HOTSCRAMBL, NEAT1 (nucleus reference gene), and GAPDH (cytoplasmic reference gene). **(G)** Representative RNA fluorescence in situ hybridization (FISH) images of *HOTSCRAMBL* in CD34^+^CD45RA^−^CD90^+^ HSPCs with either *AAVS1* knockout (control) or *HOTSCRAMBL* knockout. Scale bar, 5 µm. **(H)** Quantification of *HOTSCRAMBL*^+^ cells within the CD34^+^CD45RA^−^CD90^+^ HSPCs as determined by RNA FISH. For each replicate, 2-5 fields were analyzed (with a total of 60-120 cells counted) per group using ImageJ. All data are presented as mean ± standard deviations (SD), significance is indicated as ***P < 0.001, or n.s. (not significant).

**Supplementary Figure 2:**
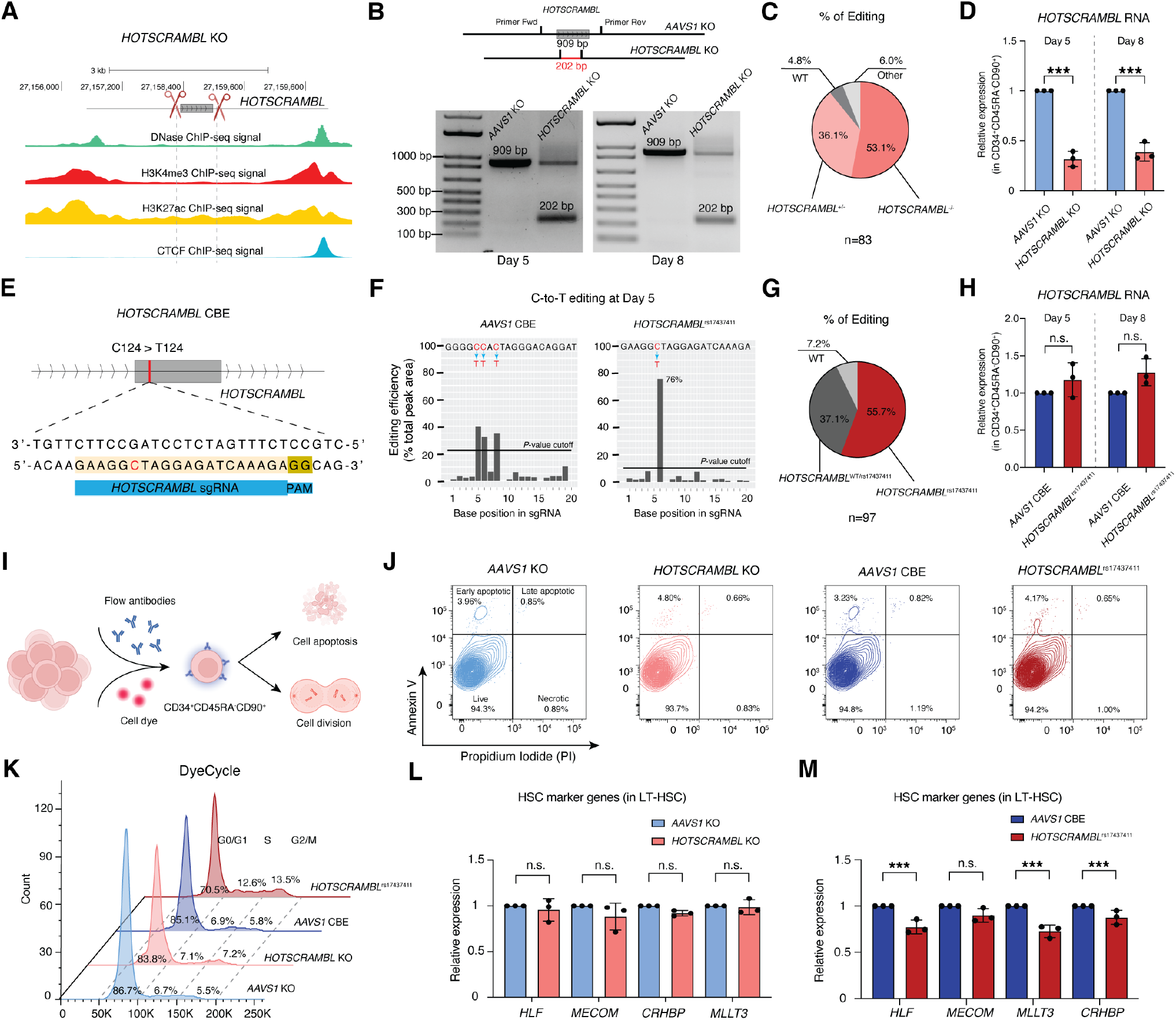
Precise base editing of *HOTSCRAMBL*^rs17437411^ in human HSPCs reveals altered transcriptional activity and HSC identity. **(A)** Schematic diagram of *HOTSCRAMBL* knockout (KO) strategy and genomic tracks from the SCREEN Registry V3^63^, showing DNase hypersensitivity and ChIP-seq signals for histone marks (H3K4me3, H3K27ac) and CTCF at the *HOTSCRAMBL* locus. **(B)** Representative gel images showing PCR products flanking the edited region of *HOTSCRAMBL* in KO groups at Day 5 and Day 8. **(C)** Percentage of HSPC genotypes at the *HOTSCRAMBL* locus shown by pie chart in KO groups at Day 5. WT (*HOTSCRAMBL*^+/+^, grey), heterozygous deletion (*HOTSCRAMBL*^+/-^, pink), homozygous deletion (*HOTSCRAMBL*^-/-^, red), or other single guides-based editing (white). **(D)** Quantification of *HOTSCRAMBL* RNA expression in CD34^+^CD45RA^−^CD90^+^ cells under *AAVS1* and *HOTSCRAMBL* KO condition at Days 5 and 8 by qPCR. Gene expression was normalized to *HPRT1*. **(E)** Schematic diagram of CBE strategy targeting the rs17437411 (C124>T124) variant site in *HOTSCRAMBL*. **(F)** Quantification of T editing frequency at each base position in *AAVS1* and *HOTSCRAMBL* sgRNA sequence at Day 5 by EditR^64^. **(G)** Pie chart showing CBE outcomes at rs17437411 in *HOTSCRAMBL* on Day 5. WT (*HOTSCRAMBL*^+/+^, grey), heterozygous editing (*HOTSCRAMBL*^rs17437411/+^, pink), homozygous editing (*HOTSCRAMBL*^rs17437411^, red). **(H)** Quantification of *HOTSCRAMBL* RNA expression in CD34^+^CD45RA^−^CD90^+^ HSPCs under *AAVS1* CBE and *HOTSCRAMBL*^rs17437411^ condition at Days 5 and 8 by qPCR. Gene expression was normalized to *HPRT1*. **(I)** Schematic overview of downstream flow cytometry analyses to assess apoptosis and cell division in CD34^+^CD45RA^−^CD90^+^ populations. **(J)**Representative flow cytometry plots showing Annexin V/PI staining of apoptotic, necrotic, and live cells in different editing conditions at Day 5. **(K)** Representative flow cytometry histograms of CD34^+^CD45RA^−^CD90^+^ cell cycle at Day 5 by DyeCycle staining. Cell cycle phase distributions are denoted as G0/G1, S and G2/M. **(L and M)** Quantification of HSC marker genes (*HLF, MECOM, CRHBP, MLLT3*) in sorted LT-HSCs by qPCR at Day 5 following *AAVS1* KO and *HOTSCRAMBL* KO (L), and *AAVS1* CBE and *HOTSCRAMBL*^rs17437411^ editing (M). Gene expression was normalized to *HPRT1*. All data are presented as mean ± SD, significance is indicated as \*\*\**P* < 0.001, or n.s. (not significant).

**Supplementary Figure 3:**
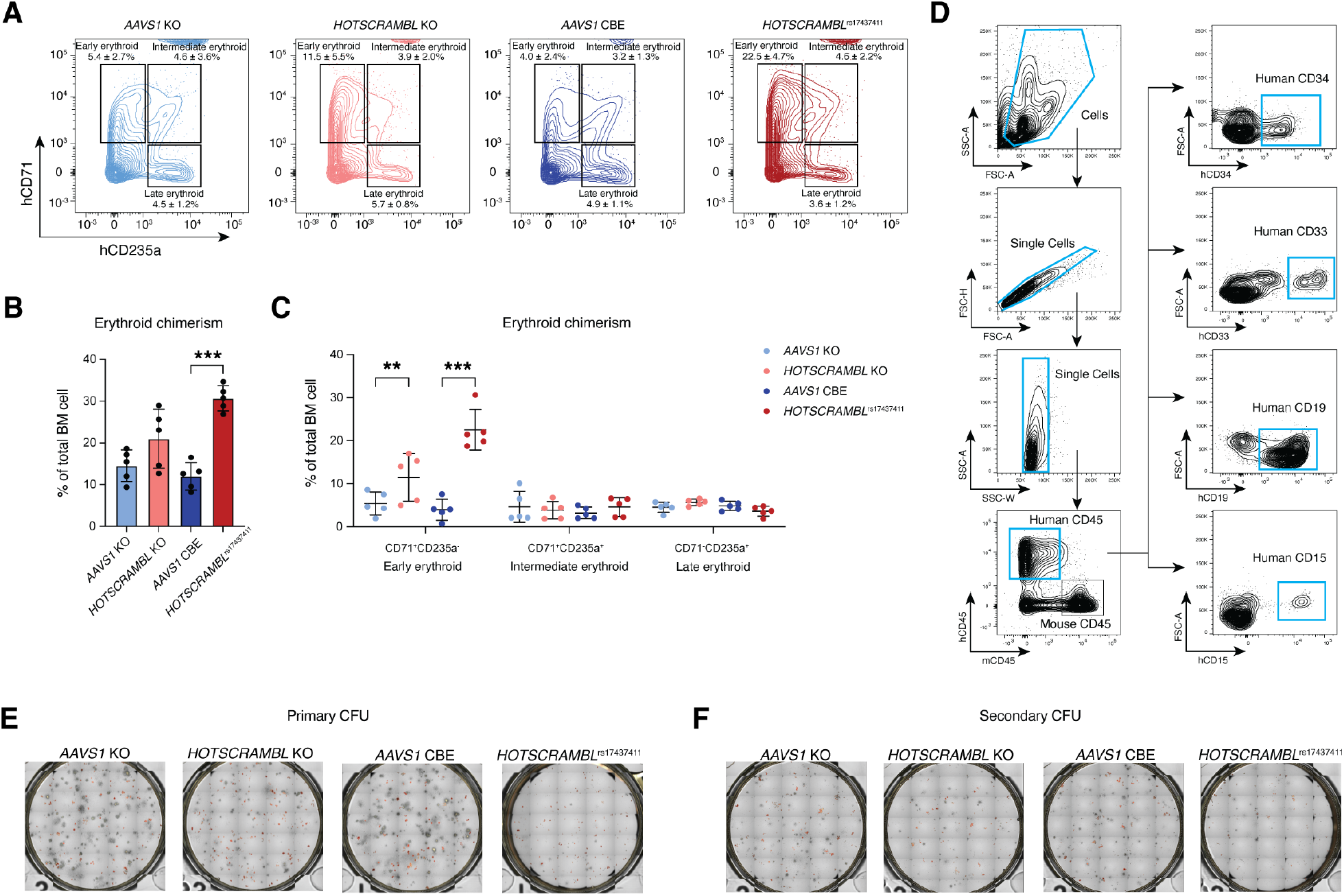
*HOTSCRAMBL*^rs17437411^ enhances early erythroid commitment and reduces clonogenic potential. **(A)** Flow cytometry gating strategy for identification of human hematopoietic populations (human CD45^+^) in BM at 16-week post transplantation, including CD34^+^ HSPCs, CD33^+^ myeloid progenitors, CD19^+^ B cells, and CD15^+^ granulocytes. **(B)** Representative flow cytometry plots of early erythroid (CD71^+^CD235a^−^), intermediate erythroid (CD71^+^CD235a^+^), and late erythroid (CD71^−^CD235a^+^) populations in BM samples at week 16 post-transplantation. **(C)** Quantification of total human erythroid chimerism in BM (% of total BM cells). **(D)** Quantification of early, intermediate, and late erythroid subpopulations based on CD71 and CD235a expression. **(E and F)** Representative plate view of methylcellulose-based CFU assays showing overall colony outputs in primary (E) and secondary (F) plating. All data are presented as mean ± SD, significance is indicated as \*\**P* < 0.01, \*\*\**P* < 0.001, or n.s. (not significant).

**Supplementary Figure 4:**
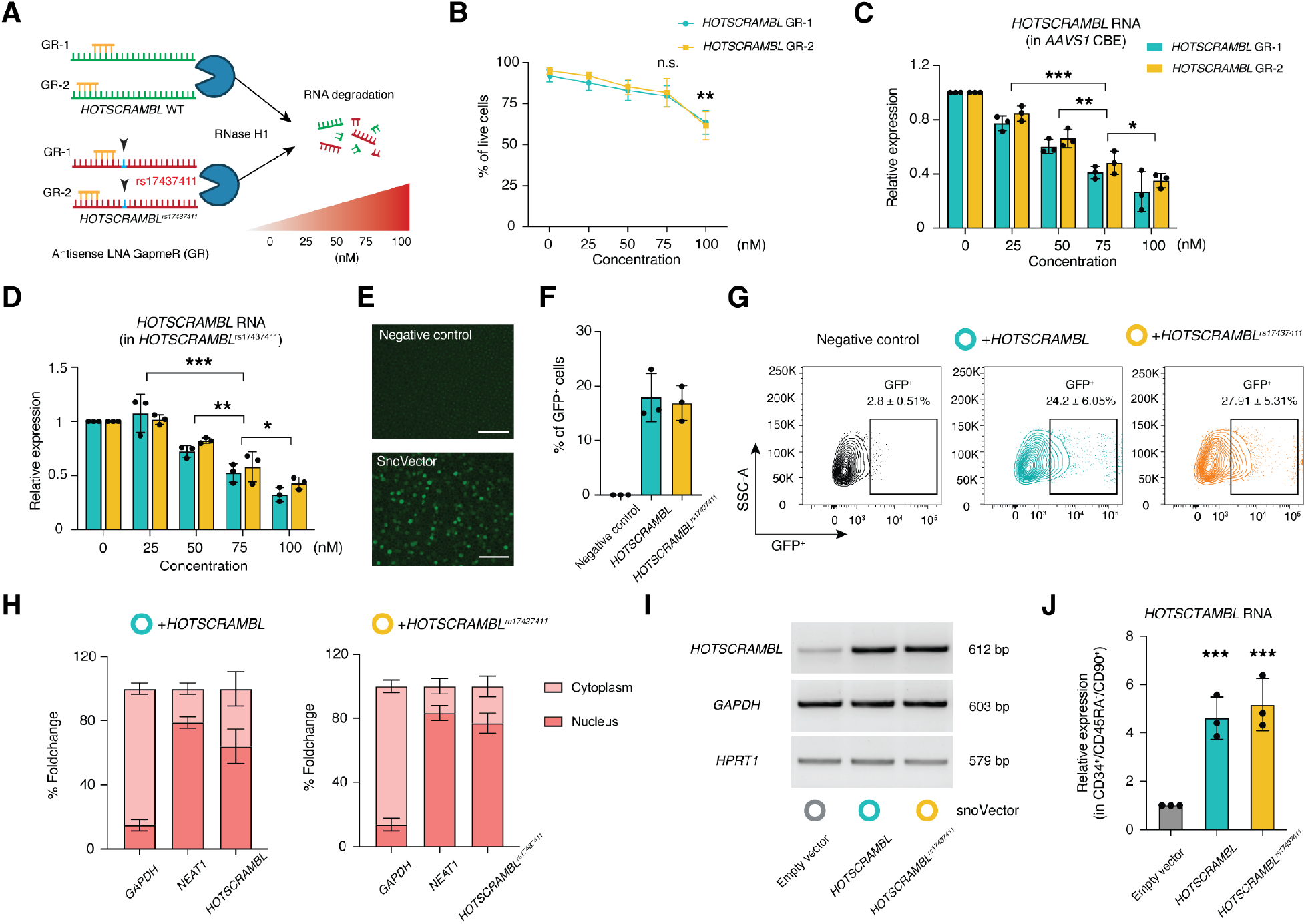
Targeted manipulation of both wild-type and *rs17437411* variant *HOTSCRAMBL* RNA expression using antisense GapmeRs and SnoVector. **(A)** Schematic of antisense LNA GapmeR-1 and GapmeR-2 target the region upstream of rs17437411, designed to degrade both *HOTSCRAMBL* and *HOTSCRAMBL*^rs17437411^ transcripts via RNase H-mediated cleavage. **(B)** Cell viability analysis following transduction with 0, 25, 50, 75, 100 nM of GapmeR-1 (GR-1) or GapmeR-2 (GR-2). **(C and D)** RNA expression levels of *HOTSCRAMBL* and *HOTSCRAMBL*^rs17437411^ after GapmeR-1 or GapmeR-2 treatment in *AAVS1* CBE (C) and *HOTSCRAMBL*^rs17437411^ (D) edited groups, respectively. Gene expression was normalized to *HPRT1*. **(E)** Fluorescence microscopy images of cells transduced with SnoVectors (bottom panel), and negative control (up panel). Scale bars, 100 µm. **(F)** Percentage of GFP^+^ cells transduced with *HOTSCRAMBL* or *HOTSCRAMBL*^rs17437411^. **(G)** Representative flow cytometry plots showing percentage of GFP^+^ cells in CD34^+^ HSPCs transduced with SnoVectors expressing *HOTSCRAMBL* or *HOTSCRAMBL*^rs17437411^. **(H)** Quantification of subcellular localization (% foldchange in nuclear/cytoplasmic fractionation) of *HOTSCRAMBL* (left panel) or *HOTSCRAMBL*^rs17437411^ (right panel), compared with nuclear-enriched *NEAT1* and cytoplasmic *GAPDH*, by qPCR following SnoVector transduction. **(I)** RT-PCR analysis showing expression of *HOTSCRAMBL, GAPDH*, and *HPRT1* in CD34^+^ HSPCs transduced with either empty vector or SnoVectors expressing *HOTSCRAMBL* or *HOTSCRAMBL*^rs17437411^, respectively. **(J)** Quantification of HOTSCRAMBL expression in CD34^+^CD45RA^−^CD90^+^ cells transduced with either empty vector or SnoVectors expressing *HOTSCRAMBL* or *HOTSCRAMBL*^rs17437411^, respectively. Gene expression was normalized to *HPRT1*. All data are presented as mean ± SD, significance is indicated as ∗*P* < 0.05, ∗∗*P* < 0.01, ∗∗∗*P* < 0.001.

**Supplementary Figure 5:**
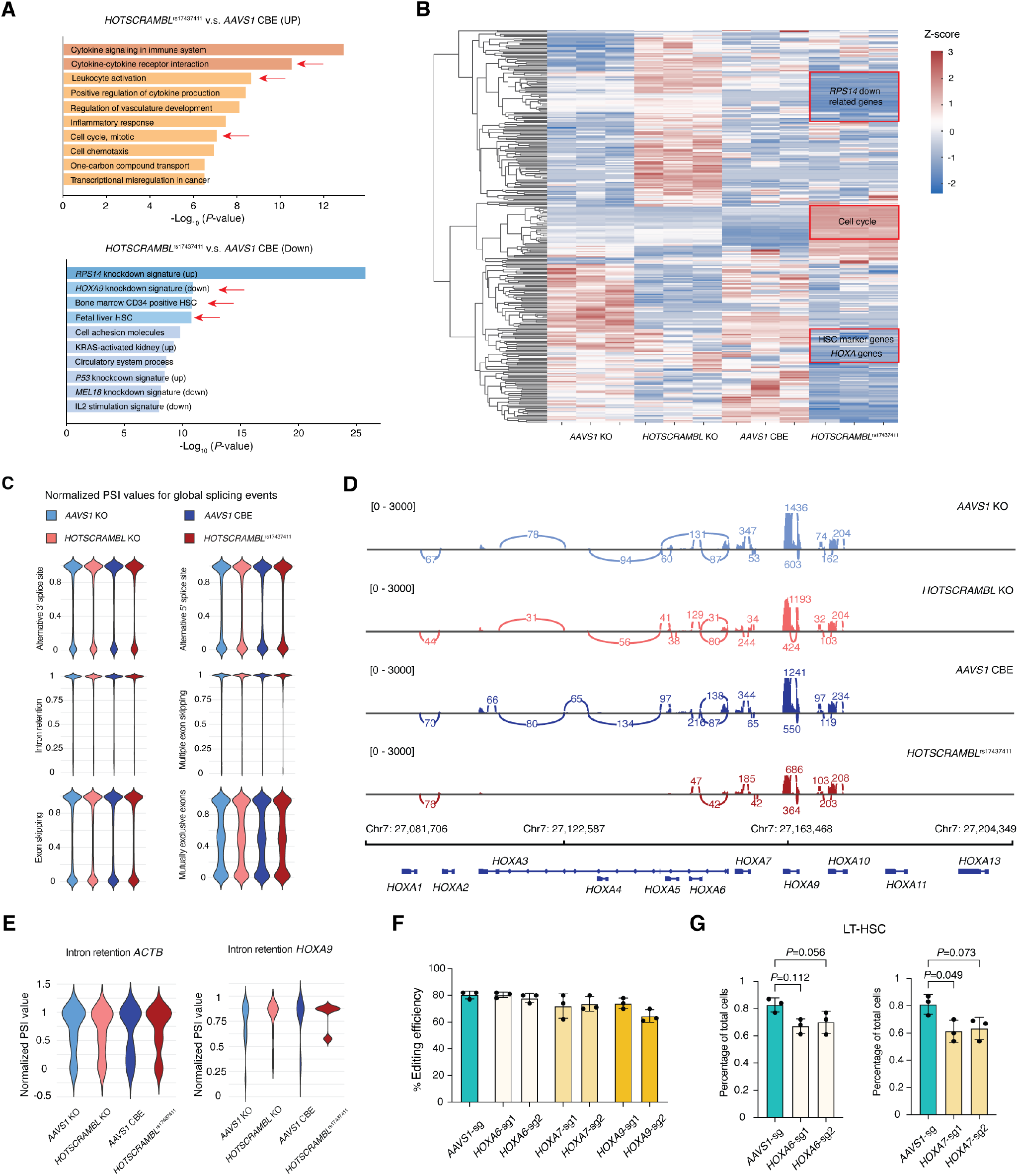
*HOTSCRAMBL*^rs17437411^ selectively disrupts *HOXA* locus splicing, identifying *HOXA9* as a major contributor to HSC defects. **(A)** Gene set enrichment analysis of up- and downregulated genes in *HOTSCRAMBL*^rs17437411^ vs *AAVS1* CBE using Metascape^65^. Top enriched terms ranked by -log_10_(*P*-value) are shown for each group, See also **Table S4. (B)** Hierarchical clustering heatmap of top DEGs *AAVS1* KO, *HOTSCRAMBL* KO, *AAVS1* CBE and *HOTSCRAMBL*^rs17437411^ conditions. Bar color represents normalized expression. **(C)** Alternative splicing analysis performed using SplAdder^66^. Violin plots show normalized percent spliced-in (PSI) values for global alternative splicing events, including alternative 3’/5’ splice sites, exon skipping, and intron retention. **(D)** Sashimi plots showing *HOXA* gene locus splicing junctions and read coverage across *AAVS1* KO, *HOTSCRAMBL* KO, *AAVS1* CBE and *HOTSCRAMBL*^rs17437411^ conditions. **(E)** Quantification of top hit intron retention events in *ACTB* (control) and *HOXA9* using SplAdder-based PSI estimates. Violin plots show normalized PSI values. **(F)** Percentage of editing efficiency (% of frameshift indels) in *HOXA6*-, *HOXA7*- and *HOXA9*-sgRNAs edited HSPCs. **(G)** Flow cytometric quantification of LT-HSC populations in *HOXA6*-, *HOXA7*-sgRNAs edited CD34^+^ HSPCs. All data are presented as mean ± SD, significance is indicated as exact *P* value.

**Supplementary Figure 6:**
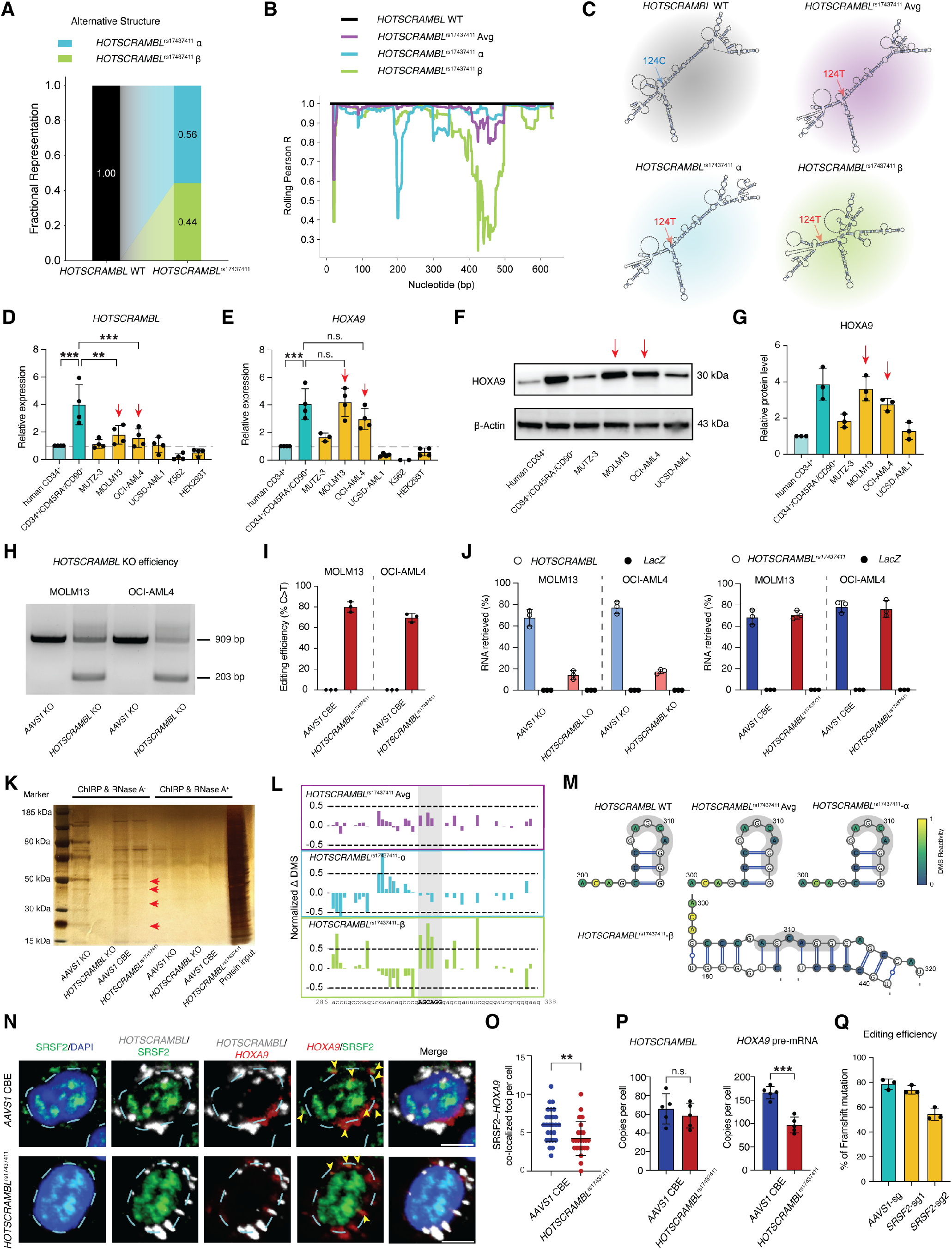
MOLM13 and OCI-AML4 serve as relevant cell models to study *HOTSCRAMBL* function, with predicted secondary structure alteration and RNA-protein binding. **(A)** Fraction of reads going to each cluster in *HOTSCRAMBL* WT and *HOTSCRAMBL*^rs17437411^ RNA. **(B)** Rolling Pearson R for each *HOTSCRAMBL*^rs17437411^ conformation relative to *HOTSCRAMBL* WT RNA. **(C)** Predicted structures with DMS constraints. Red arrows denote rs17437411 variant. **(D and E)** Expression of *HOTSCRAMBL* (D) and *HOXA9* (E) in cell lines, including MUTZ-3, MOLM13, OCI-AML4, UCSD-AML1, K562 and HEK293T. Gene expression was normalized to *HPRT1* and compared to CD34^+^ HSPCs as a reference. Red arrows indicate cell lines with relatively high expression of indicated genes. **(F)** Western blot of HOXA9 protein levels in AML cell lines, including MUTZ-3, MOLM13, OCI-AML4, UCSD-AML1. CD34^+^ HSPCs used as reference. β-Actin used as loading control. Cell lines with high HOXA9 level are denoted in red arrows. **(G)** Quantification of western blot in (F) by imageJ, Cell lines with high HOXA9 level are denoted in red arrows. **(H)** Agarose gel electrophoresis showing *HOTSCRAMBL* knockout efficiency in MOLM13 and OCI-AML4 cells. PCR amplicons of *HOTSCRAMBL* locus in *AAVS1* KO (909 bp) and *HOTSCRAMBL* KO (203 bp) are indicated. **(I)** CBE efficiency (% of C>T conversion) at *HOTSCRAMBL*^rs17437411^ locus in MOLM13 and OCI-AML4 cells. **(J)** Percentage of RNA pull-down for *HOTSCRAMBL* and *LacZ* control transcripts in MOLM13 and OCI-AML4 cells using biotinylated oligos. **(K)** Silver-stained gel of ChIRP-MS eluates under RNase A^-^ or RNase A^+^ conditions. Red arrows indicate differential enriched protein bands. **(L)** Normalized change in DMS signal (*HOTSCRAMBL* WT minus [*HOTSCRAMBL*^rs17437411^-Avg (purple peaks), *HOTSCRAMBL*^rs17437411^-α (cyan peaks) and *HOTSCRAMBL*^rs17437411^-β (green peaks]) around the 308 SRSF2 binding site (grey shaded area). **(M)** Predicted RNA structure at the SRSF2 308-binding site colored by DMS reactivity. **(N)** Representative RNA FISH and immunofluorescence images showing co-localization of *HOTSCRAMBL* (white), *HOXA9* pre-mRNA (red), and SRSF2 (green) in control (*AAVS1*-CBE) and *HOTSCRAMBL*^rs17437411^-edited HSPCs. Nuclei (DAPI) are outlined with dashed blue circles. Co-localization of SRSF2 and *HOXA9* is indicated by yellow arrows. Scale bar, 5 μm. **(O)** Quantification of SRSF2-*HOXA9* co-localization per cell as determined by fluorescence imaging. **(P)** Absolute qPCR quantification of *HOTSCRAMBL* or *HOTSCRAMBL*^rs17437411^, and *HOXA9* pre-mRNA copy number per cell in MOLM13 cells with *AAVS1* CBE or *HOTSCRAMBL*^rs17437411^ editing. **(Q)** Bar graph showing editing efficiency (% of frameshift mutation) in MOLM3 cells using guideRNA *SRSF2*-sg1 or *SRSF2*-sg2. All data are presented as mean ± SD, significance is indicated as \**P* < 0.05, \*\**P* < 0.01, \*\*\**P* < 0.001, or n.s. (not significant).

**Supplementary Figure 7:**
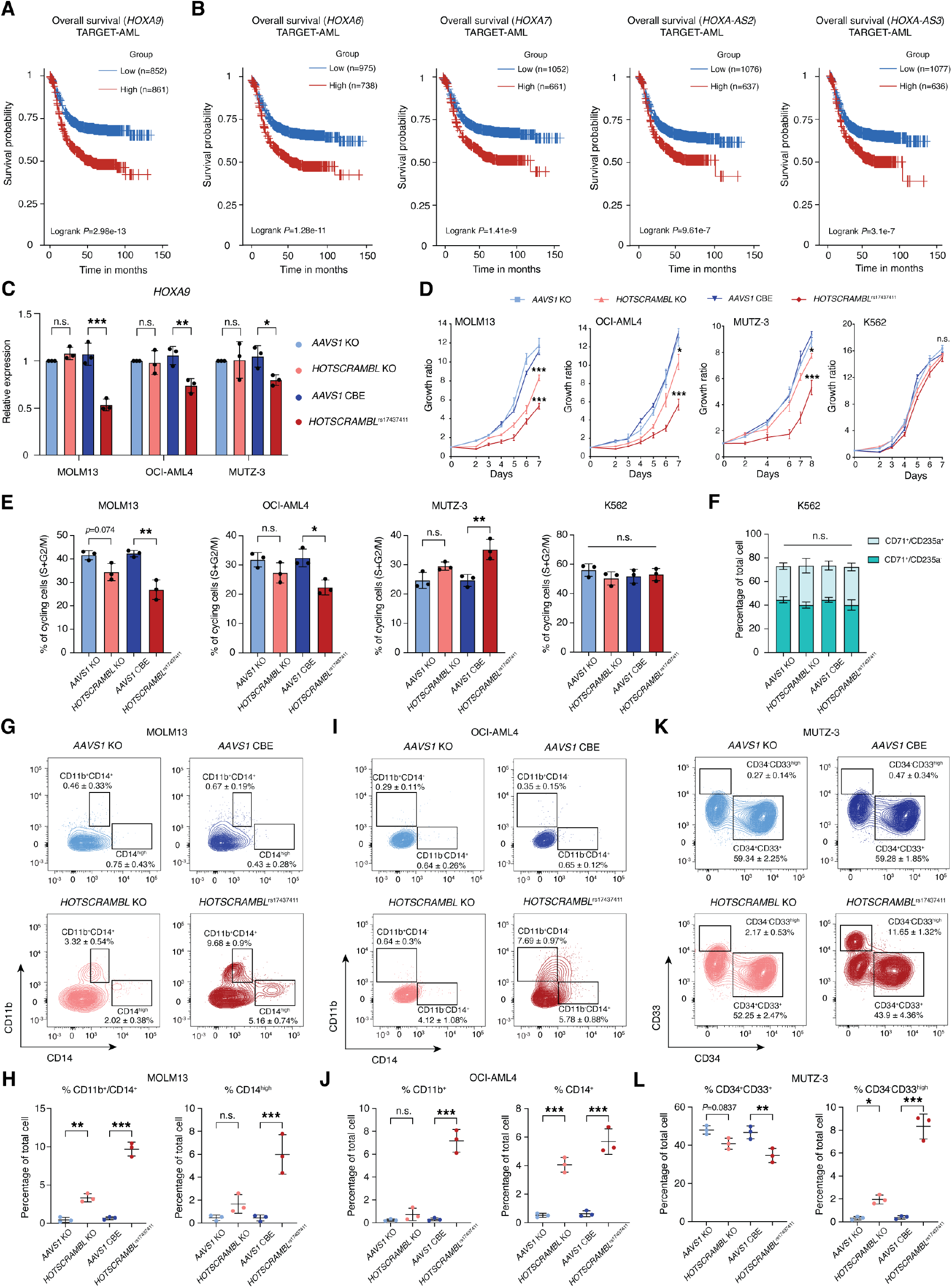
High *HOXA* genes are highly associated with poor prognosis in AML and *HOTSCRAMBL* variant selectively impairs leukemic cell growth and differentiation. **(A)** Kaplan-Meier overall survival curve of pediatric AML patients in the TARGET-AML cohort stratified by *HOXA9* expression levels (Low, n = 852; High, n = 861). **(B)** Kaplan-Meier survival curves from the TARGET-AML dataset comparing patients with high (red) and low (blue) expression of *HOXA6* (left), *HOXA7* (center-left), *HOXA*-*AS2* (center-right), and *HOXA*-*AS3* (right). **(C)** Quantification of *HOXA9* mRNA expression in edited AML cell lines (MOLM13, OCI-AML4, and MUTZ-3) by qPCR. **(D)** Cell growth ratio of MOLM13, OCI-AML4, MUTZ-3, and K562 cells measured for 7 or 8 days post-plating. **(E)** Quantification of the percentage of cells in S/G2/M phases at Day 4 post-editing in MOLM13, OCI-AML4 MUTZ-3, and K562 cells, as measured by DyeCycle staining. **(F)** Quantification of the percentage of CD71^+^CD235a^−^ and CD71^+^CD235a^+^ cells in K562 at Day 4 post-editing. **(G)** Representative flow cytometric plots of the percentage of CD11b^+^CD14^+^ and CD14^high^ cells in MOLM13 at Day 4 post-editing. **(H)** Representative flow cytometric plots of the percentage of CD11b^+^ and CD14^+^ cells in OCI-AML4 at Day 4 post-editing. **(I)** Representative flow cytometric plots of the percentage of CD34^+^CD33^+^ and CD34^-^CD33^high^ cells in MUTZ-3 at Day 4 post-editing. **(J)** Flow cytometric quantification of the percentage of CD11b^+^CD14^+^ and CD14^high^ cells in (G). **(K)** Flow cytometric quantification of the percentage of CD11b^+^ and CD14^+^ cells in (H). **(L)** Flow cytometric quantification of the percentage of CD34^+^CD33^+^ and CD34^-^CD33^high^ cells in (I). All data are presented as mean ± SD, significance is indicated as \**P* < 0.05, \*\**P* < 0.01, \*\*\**P* < 0.001, or n.s. (not significant).

## Notes

### Summary of Updates

Revised manuscript with additional experiments and updated figures; updated analysis and improved clarity of the text; updated author list and contribution; supplemental files updated.

## References

1. Afzal, Z., and Krumlauf, R. (2022). Transcriptional Regulation and Implications for Controlling Hox Gene Expression. Journal of developmental biology 10. 10.3390/jdb10010004.

2. Deschamps, J., and Duboule, D. (2017). Embryonic timing, axial stem cells, chromatin dynamics, and the Hox clock. Genes Dev 31, 1406– 1416.

3. Pearson, J.C., Lemons, D., and McGinnis, W. (2005). Modulating Hox gene functions during animal body patterning. Nat Rev Genet 6, 893– 904.

4. Krumlauf, R. (1994). Hox genes in vertebrate development. Cell 78. 10.1016/0092-8674(94)90290-9.

5. Fowler, J.L., Zheng, S.L., Nguyen, A., Chen, A., Xiong, X., Chai, T., Chen, J.Y., Karigane, D., Banuelos, A.M., Niizuma, K., et al. (2024). Lineage-tracing hematopoietic stem cell origins in vivo to efficiently make human HLF+ HOXA+ hematopoietic progenitors from pluripotent stem cells. Dev Cell 59, 1110–1131.e22.

6. Bach, C., Buhl, S., Mueller, D., García-Cuéllar, M.-P., Maethner, E., and Slany, R.K. (2010). Leukemogenic transformation by HOXA cluster genes. Blood 115, 2910–2918.

7. Dou, D.R., Calvanese, V., Sierra, M.I., Nguyen, A.T., Minasian, A., Saarikoski, P., Sasidharan, R., Ramirez, C.M., Zack, J.A., Crooks, G.M., et al. (2016). Medial HOXA genes demarcate haematopoietic stem cell fate during human development. Nature cell biology 18. 10.1038/ncb3354.

8. Luo, H., Zhu, G., Xu, J., Lai, Q., Yan, B., Guo, Y., Fung, T.K., Zeisig, B.B., Cui, Y., Zha, J., et al. (2019). HOTTIP lncRNA Promotes Hematopoietic Stem Cell Self-Renewal Leading to AML-like Disease in Mice. Cancer Cell 36, 645–659.e8.

9. Collins, C.T., and Hess, J.L. (2016). Role of HOXA9 in leukemia: dysregulation, cofactors and essential targets. Oncogene 35, 1090– 1098.

10. Sun, Y., Zhou, B., Mao, F., Xu, J., Miao, H., Zou, Z., Phuc Khoa, L.T., Jang, Y., Cai, S., Witkin, M., et al. (2018). HOXA9 Reprograms the Enhancer Landscape to Promote Leukemogenesis. Cancer Cell 34, 643– 658.e5.

11. Jung, H.S., Uenishi, G., Park, M.A., Liu, P., Suknuntha, K., Raymond, M., Choi, Y.J., Thomson, J.A., Ong, I.M., and Slukvin, I.I. (2021). SOX17 integrates HOXA and arterial programs in hemogenic endothelium to drive definitive lympho-myeloid hematopoiesis. Cell Rep 34, 108758.

12. Khan, I., Amin, M.A., Eklund, E.A., and Gartel, A.L. (2024). Regulation of HOX gene expression in AML. Blood Cancer J 14, 42.

13. Zhao, J., Cato, L.D., Arora, U.P., Bao, E.L., Bryant, S.C., Williams, N., Jia, Y., Goldman, S.R., Nangalia, J., Erb, M.A., et al. (2024). Inherited blood cancer predisposition through altered transcription elongation. Cell 187, 642–658.e19.

14. David Wang, X.Q., Gore, H., Himadewi, P., Feng, F., Yang, L., Zhou, W., Liu, Y., Wang, X., Chen, C.-W., Su, J., et al. (2020). Three-dimensional regulation ofHOXAcluster genes by acis-element in hematopoietic stem cell and leukemia. bioRxiv. 10.1101/2020.04.16.017533.

15. Luo, H., Wang, F., Zha, J., Li, H., Yan, B., Du, Q., Yang, F., Sobh, A., Vulpe, C., Drusbosky, L., et al. (2018). CTCF boundary remodels chromatin domain and drives aberrant gene transcription in acute myeloid leukemia. Blood 132, 837–848.

16. Qiu, Y., Xu, M., and Huang, S. (2021). Long noncoding RNAs: emerging regulators of normal and malignant hematopoiesis. Blood 138, 2327–2336.

17. Chen, L.-L., and Kim, V.N. (2024). Small and long non-coding RNAs: Past, present, and future. Cell 187, 6451–6485.

18. Statello, L., Guo, C.-J., Chen, L.-L., and Huarte, M. (2021). Gene regulation by long non-coding RNAs and its biological functions. Nat Rev Mol Cell Biol 22, 96–118.

19. Yin, Y., Yan, P., Lu, J., Song, G., Zhu, Y., Li, Z., Zhao, Y., Shen, B., Huang, X., Zhu, H., et al. (2015). Opposing Roles for the lncRNA Haunt and Its Genomic Locus in Regulating HOXA Gene Activation during Embryonic Stem Cell Differentiation. Cell Stem Cell 16, 504– 516.

20. Wang, K.C., Yang, Y.W., Liu, B., Sanyal, A., Corces-Zimmerman, R., Chen, Y., Lajoie, B.R., Protacio, A., Flynn, R.A., Gupta, R.A., et al. (2011). A long noncoding RNA maintains active chromatin to coordinate homeotic gene expression. Nature 472, 120–124.

21. Zhang, X., Lian, Z., Padden, C., Gerstein, M.B., Rozowsky, J., Snyder, M., Gingeras, T.R., Kapranov, P., Weissman, S.M., and Newburger, P.E. (2009). A myelopoiesis-associated regulatory intergenic noncoding RNA transcript within the human HOXA cluster. Blood 113, 2526–2534.

22. Wang, X.Q.D., and Dostie, J. (2017). Reciprocal regulation of chromatin state and architecture by HOTAIRM1 contributes to temporal collinear HOXA gene activation. Nucleic Acids Res 45, 1091–1104.

23. Bartell, E., Lin, K., Tsuo, K., Gan, W., Vedantam, S., Cole, J.B., Baronas, J.M., Yengo, L., Marouli, E., Amariuta, T., et al. (2026). Genetics of skeletal proportions across two different populations. Am J Hum Genet 113, 794–808.

24. Bao, E.L., Nandakumar, S.K., Liao, X., Bick, A.G., Karjalainen, J., Tabaka, M., Gan, O.I., Havulinna, A.S., Kiiskinen, T.T.J., Lareau, C.A., et al. (2020). Inherited myeloproliferative neoplasm risk affects haematopoietic stem cells. Nature 586, 769–775.

25. Werner, B., Beier, F., Hummel, S., Balabanov, S., Lassay, L., Orlikowsky, T., Dingli, D., Brümmendorf, T.H., and Traulsen, A. (2015). Reconstructing the in vivo dynamics of hematopoietic stem cells from telomere length distributions. 10.7554/eLife.08687.

26. Brown, D.W., Cato, L.D., Zhao, Y., Nandakumar, S.K., Bao, E.L., Gardner, E.J., Hubbard, A.K., DePaulis, A., Rehling, T., Song, L., et al. (2023). Shared and distinct genetic etiologies for different types of clonal hematopoiesis. Nature Communications 14, 1–13.

27. Bruhn-Olszewska, B., Markljung, E., Rychlicka-Buniowska, E., Sarkisyan, D., Filipowicz, N., and Dumanski, J.P. (2025). The effects of loss of Y chromosome on male health. Nature Reviews Genetics 26, 320– 335.

28. Voit, R.A., Tao, L., Yu, F., Cato, L.D., Cohen, B., Fleming, T.J., Antoszewski, M., Liao, X., Fiorini, C., Nandakumar, S.K., et al. (2022). A genetic disorder reveals a hematopoietic stem cell regulatory network co-opted in leukemia. Nature Immunology 24, 69–83.

29. Yin, Q.-F., Hu, S.-B., Xu, Y.-F., Yang, L., Carmichael, G.G., and Chen, L.-L. (2015). SnoVectors for nuclear expression of RNA. Nucleic Acids Res 43, e5.

30. Romero-Barrios, N., Legascue, M.F., Benhamed, M., Ariel, F., and Crespi, M. (2018). Splicing regulation by long noncoding RNAs. Nucleic Acids Res 46, 2169–2184.

31. Nojima, T., Rebelo, K., Gomes, T., Grosso, A.R., Proudfoot, N.J., and Carmo-Fonseca, M. (2018). RNA Polymerase II Phosphorylated on CTD Serine 5 Interacts with the Spliceosome during Co-transcriptional Splicing. Mol Cell 72, 369–379.e4.

32. Engreitz, J.M., Haines, J.E., Perez, E.M., Munson, G., Chen, J., Kane, M., McDonel, P.E., Guttman, M., and Lander, E.S. (2016). Local regulation of gene expression by lncRNA promoters, transcription and splicing. Nature 539, 452–455.

33. Wenge, D.V., and Armstrong, S.A. (2024). The future of HOXAexpressing leukemias: Menin inhibitor response and resistance. Current opinion in hematology 31. 10.1097/MOH.0000000000000796.

34. Zubradt, M., Gupta, P., Persad, S., Lambowitz, A.M., Weissman, J.S., and Rouskin, S. (2017). DMS-MaPseq for genome-wide or targeted RNA structure probing in vivo. Nat Methods 14, 75–82.

35. Chu, C., Qu, K., Zhong, F.L., Artandi, S.E., and Chang, H.Y. (2011). Genomic maps of long noncoding RNA occupancy reveal principles of RNA-chromatin interactions. Molecular cell 44. 10.1016/j.molcel.2011.08.027.

36. Chu, C., and Chang, H.Y. (2018). ChIRP-MS: RNA-Directed Proteomic Discovery. Methods Mol Biol 1861, 37–45.

37. Paz, I., Kosti, I., Ares, M., Jr, Cline, M., and Mandel-Gutfreund, Y. (2014). RBPmap: a web server for mapping binding sites of RNA-binding proteins. Nucleic Acids Res 42, W361–W367.

38. Zeng, A.G.X., Bansal, S., Jin, L., Mitchell, A., Chen, W.C., Abbas, H.A., Chan-Seng-Yue, M., Voisin, V., van Galen, P., Tierens, A., et al. (2022). A cellular hierarchy framework for understanding heterogeneity and predicting drug response in acute myeloid leukemia. Nature Medicine 28, 1212–1223.

39. Shah, N., and Sukumar, S. (2010). The Hox genes and their roles in oncogenesis. Nat Rev Cancer 10, 361–371.

40. Hubert, K.A., and Wellik, D.M. (2023). Hox genes in development and beyond. Development 150. 10.1242/dev.192476.

41. Gupta, R.A., Shah, N., Wang, K.C., Kim, J., Horlings, H.M., Wong, D.J., Tsai, M.-C., Hung, T., Argani, P., Rinn, J.L., et al. (2010). Long non-coding RNA HOTAIR reprograms chromatin state to promote cancer metastasis. Nature 464, 1071–1076.

42. Qin, Y., Ren, J., Yu, H., He, X., Cheng, S., Chen, W., Yang, Z., Sun, F., Wang, C., Yuan, S., et al. (2024). HOXA-AS2 Epigenetically Inhibits HBV Transcription by Recruiting the MTA1-HDAC1/2 Deacetylase Complex to cccDNA Minichromosome. Adv Sci (Weinh) 11, e2306810.

43. Degani, N., Lubelsky, Y., Perry, R.B.-T., Ainbinder, E., and Ulitsky, I. (2021). Highly conserved and cis-acting lncRNAs produced from paralogous regions in the center of HOXA and HOXB clusters in the endoderm lineage. PLoS Genet 17, e1009681.

44. Isaev, K., Jiang, L., Wu, S., Lee, C.A., Watters, V., Fort, V., Tsai, R., Coutinho, F.J., Hussein, S.M.I., Zhang, J., et al. (2021). Pan-cancer analysis of non-coding transcripts reveals the prognostic onco-lncRNA HOXA10-AS in gliomas. Cell Rep 37, 109873.

45. Kim, E., Ilagan, J.O., Liang, Y., Daubner, G.M., Lee, S.C.-W., Ramakrishnan, A., Li, Y., Chung, Y.R., Micol, J.-B., Murphy, M.E., et al. (2015). SRSF2 Mutations Contribute to Myelodysplasia by Mutant-Specific Effects on Exon Recognition. Cancer Cell 27, 617–630.

46. Wang, B.A., Mehta, H.M., Penumutchu, S.R., Tolbert, B.S., Cheng, C., Kimmel, M., Haferlach, T., Maciejewski, J.P., and Corey, S.J. (2022). Alternatively spliced CSF3R isoforms in SRSF2 P95H mutated myeloid neoplasms. Leukemia 36, 2499–2508.

47. Liu, X., Devadiga, S.A., Stanley, R.F., Morrow, R.M., Janssen, K.A., Quesnel-Vallières, M., Pomp, O., Moverley, A.A., Li, C., Skuli, N., et al. (2024). A mitochondrial surveillance mechanism activated by SRSF2 mutations in hematologic malignancies. J Clin Invest 134. 10.1172/JCI175619.

48. Luo, H., Zhu, G., Eshelman, M.A., Fung, T.K., Lai, Q., Wang, F., Zeisig, B.B., Lesperance, J., Ma, X., Chen, S., et al. (2022). HOTTIP-de-pendent R-loop formation regulates CTCF boundary activity and TAD integrity in leukemia. Mol Cell 82, 833–851.e11.

49. Thompson, D.J., Genovese, G., Halvardson, J., Ulirsch, J.C., Wright, D.J., Terao, C., Davidsson, O.B., Day, F.R., Sulem, P., Jiang, Y., et al. (2019). Genetic predisposition to mosaic Y chromosome loss in blood. Nature 575, 652–657.

50. Agarwal, G., Antoszewski, M., Xie, X., Pershad, Y., Arora, U.P., Poon, C.-L., Lyu, P., Lee, A.J., Guo, C.-J., Ye, T., et al. (2025). Inherited resilience to clonal hematopoiesis by modifying stem cell RNA regulation. bioRxiv. 10.1101/2025.03.24.645017.

51. Graham, A., Papalopulu, N., and Krumlauf, R. (1989). The murine and Drosophila homeobox gene complexes have common features of organization and expression. Cell 57. 10.1016/00928674(89)90912-4.

52. Weng, C., Yu, F., Yang, D., Poeschla, M., Liggett, L.A., Jones, M.G., Qiu, X., Wahlster, L., Caulier, A., Hussmann, J.A., et al. (2024). Deciphering cell states and genealogies of human haematopoiesis. Nature 627. 10.1038/s41586-024-07066-z.

53. Mitchell, E., Spencer, C.M., Williams, N., Dawson, K.J., Mende, N., Calderbank, E.F., Jung, H., Mitchell, T., Coorens, T.H.H., Spencer, D.H., et al. (2022). Clonal dynamics of haematopoiesis across the human lifespan. Nature 606. 10.1038/s41586-022-04786-y.

54. Lai, F., Damle, S.S., Ling, K.K., and Rigo, F. (2020). Directed RNase H Cleavage of Nascent Transcripts Causes Transcription Termination. Mol Cell 77, 1032–1043.e4.

55. Crooke, S.T., Baker, B.F., Crooke, R.M., and Liang, X.-H. (2021). Antisense technology: an overview and prospectus. Nat Rev Drug Discov 20, 427–453.

56. Kishore, S., Khanna, A., and Stamm, S. (2008). Rapid generation of splicing reporters with pSpliceExpress. Gene 427, 104–110.

57. Dwivedi, B., Mumme, H., Satpathy, S., Bhasin, S.S., and Bhasin, M. (2022). Survival Genie, a web platform for survival analysis across pediatric and adult cancers. Sci Rep 12, 3069.

58. Kessler, M.D., Damask, A., O’Keeffe, S., Banerjee, N., Li, D., Watanabe, K., Marketta, A., Van Meter, M., Semrau, S., Horowitz, J., et al. (2022). Common and rare variant associations with clonal haematopoiesis phenotypes. Nature 612, 301–309.

59. Jaganathan, K., Kyriazopoulou Panagiotopoulou, S., McRae, J.F., Darbandi, S.F., Knowles, D., Li, Y.I., Kosmicki, J.A., Arbelaez, J., Cui, W., Schwartz, G.B., et al. (2019). Predicting Splicing from Primary Sequence with Deep Learning. Cell 176, 535–548.e24.

60. Zeng, T., and Li, Y.I. (2022). Predicting RNA splicing from DNA sequence using Pangolin. Genome Biol 23, 103.

61. Murphy, W.J., Eizirik, E., O’Brien, S.J., Madsen, O., Scally, M., Douady, C.J., Teeling, E., Ryder, O.A., Stanhope, M.J., de Jong, W.W., et al. (2001). Resolution of the early placental mammal radiation using Bayesian phylogenetics. Science 294, 2348–2351.

62. Li, Z., Liu, L., Feng, C., Qin, Y., Xiao, J., Zhang, Z., and Ma, L. (2023). LncBook 2.0: integrating human long non-coding RNAs with multiomics annotations. Nucleic Acids Res 51, D186–D191.

63. ENCODE Project Consortium, Moore, J.E., Purcaro, M.J., Pratt, H.E., Epstein, C.B., Shoresh, N., Adrian, J., Kawli, T., Davis, C.A., Dobin, A., et al. (2020). Expanded encyclopaedias of DNA elements in the human and mouse genomes. Nature 583, 699–710.

64. Kluesner, M.G., Nedveck, D.A., Lahr, W.S., Garbe, J.R., Abrahante, J.E., Webber, B.R., and Moriarity, B.S. (2018). EditR: A Method to Quantify Base Editing from Sanger Sequencing. CRISPR J 1, 239–250.

65. Zhou, Y., Zhou, B., Pache, L., Chang, M., Khodabakhshi, A.H., Tanaseichuk, O., Benner, C., and Chanda, S.K. (2019). Metascape provides a biologist-oriented resource for the analysis of systems-level datasets. Nat Commun 10, 1523.

66. Kahles, A., Ong, C.S., Zhong, Y., and Rätsch, G. (2016). SplAdder: identification, quantification and testing of alternative splicing events from RNA-Seq data. Bioinformatics 32, 1840–1847.

67. Matsuo, Y., MacLeod, R.A., Uphoff, C.C., Drexler, H.G., Nishizaki, C., Katayama, Y., Kimura, G., Fujii, N., Omoto, E., Harada, M., et al. (1997). Two acute monocytic leukemia (AML-M5a) cell lines (MOLM13 and MOLM-14) with interclonal phenotypic heterogeneity showing MLL-AF9 fusion resulting from an occult chromosome insertion, ins(11;9)(q23;p22p23). Leukemia 11, 1469–1477.

68. Koistinen, P., Wang, C., Yang, G.S., Wang, Y.F., Williams, D.E., Lyman, S.D., Minden, M.D., and McCulloch, E.A. (1991). OCI/AML-4 an acute myeloblastic leukemia cell line: regulation and response to cytosine arabinoside. Leukemia 5, 704–711.

69. Drexler, H.G., Zaborski, M., and Quentmeier, H. (1997). Cytokine response profiles of human myeloid factor-dependent leukemia cell lines. Leukemia 11, 701–708.

70. Townsend, E.C., Murakami, M.A., Christodoulou, A., Christie, A.L., Köster, J., DeSouza, T.A., Morgan, E.A., Kallgren, S.P., Liu, H., Wu, S.-C., et al. (2016). The Public Repository of Xenografts Enables Discovery and Randomized Phase II-like Trials in Mice. Cancer Cell 29, 574– 586.

71. Zhao, J., Jia, Y., Mahmut, D., Deik, A.A., Jeanfavre, S., Clish, C.B., and Sankaran, V.G. (2023). Human hematopoietic stem cell vulnerability to ferroptosis. Cell 186, 732–747.e16.

72. Vuckovic, D., Bao, E.L., Akbari, P., Lareau, C.A., Mousas, A., Jiang, T., Chen, M.-H., Raffield, L.M., Tardaguila, M., Huffman, J.E., et al. (2020). The Polygenic and Monogenic Basis of Blood Traits and Diseases. Cell 182, 1214–1231.e11.

73. Howrigan, D.P., Abbott, L., rkwalters, Palmer, D., Francioli, L., and Hammerbacher, J. (2023). Nealelab/UK_Biobank_GWAS: v2 (Zenodo) 10.5281/ZENODO.8011557.

74. Codd, V., Denniff, M., Swinfield, C., Warner, S.C., Papakonstantinou, M., Sheth, S., Nanus, D.E., Budgeon, C.A., Musicha, C., Bountziouka, V., et al. (2022). Measurement and initial characterization of leukocyte telomere length in 474,074 participants in UK Biobank. Nature aging 2. 10.1038/s43587-021-00166-9.

75. Poeschla, M., Arora, U.P., Walne, A., McReynolds, L.J., Niewisch, M.R., Giri, N., Zeigler, L.P., Gusev, A., Machiela, M.J., Tummala, H., et al. (2025). Polygenic modifiers impact penetrance and expressivity in telomere biology disorders. J Clin Invest. 10.1172/JCI191107.

76. Weissbrod, O., Hormozdiari, F., Benner, C., Cui, R., Ulirsch, J., Gazal, S., Schoech, A.P., van de Geijn, B., Reshef, Y., Márquez-Luna, C., et al. (2020). Functionally informed fine-mapping and polygenic localization of complex trait heritability. Nat Genet 52, 1355–1363.

77. Wang, G., Sarkar, A., Carbonetto, P., and Stephens, M. (2020). A simple new approach to variable selection in regression, with application to genetic fine mapping. J R Stat Soc Series B Stat Methodol 82, 1273– 1300.

78. Yang, Z., Wang, C., Liu, L., Khan, A., Lee, A., Vardarajan, B., Mayeux, R., Kiryluk, K., and Ionita-Laza, I. (2023). CARMA is a new Bayesian model for fine-mapping in genome-wide association meta-analyses. Nat Genet 55, 1057–1065.

79. Martin-Rufino, J.D., Castano, N., Pang, M., Grody, E.I., Joubran, S., Caulier, A., Wahlster, L., Li, T., Qiu, X., Riera-Escandell, A.M., et al. (2023). Massively parallel base editing to map variant effects in human hematopoiesis. Cell 186, 2456–2474.e24.

80. Martin-Rufino, J.D., Caulier, A., Lee, S., Castano, N., King, E., Joubran, S., Jones, M., Goldman, S.R., Arora, U.P., Wahlster, L., et al. (2025). Transcription factor networks disproportionately enrich for heritability of blood cell phenotypes. Science 388, 52–59.

81. Arbab, M., Shen, M.W., Mok, B., Wilson, C., Matuszek, Ż., Cassa, C.A., and Liu, D.R. (2020). Determinants of Base Editing Outcomes from Target Library Analysis and Machine Learning. Cell 182, 463–480.e30.

82. Hill, J.T., Demarest, B.L., Bisgrove, B.W., Su, Y.-C., Smith, M., and Yost, H.J. (2014). Poly peak parser: Method and software for identification of unknown indels using sanger sequencing of polymerase chain reaction products. Dev Dyn 243, 1632–1636.

83. Brinkman, E.K., Chen, T., Amendola, M., and van Steensel, B. (2014). Easy quantitative assessment of genome editing by sequence trace decomposition. Nucleic Acids Res 42, e168.

84. Chen, L.-L., DeCerbo, J.N., and Carmichael, G.G. (2008). Alu elementmediated gene silencing. EMBO J 27, 1694–1705.

85. Tripathi, V., Fei, J., Ha, T., and Prasanth, K.V. (2015). RNA fluorescence in situ hybridization in cultured mammalian cells. Methods Mol Biol 1206, 123–136.

86. Guo, C.-J., Ma, X.-K., Xing, Y.-H., Zheng, C.-C., Xu, Y.-F., Shan, L., Zhang, J., Wang, S., Wang, Y., Carmichael, G.G., et al. (2020). Distinct Processing of lncRNAs Contributes to Non-conserved Functions in Stem Cells. Cell 181, 621–636.e22.

87. Chen, K.H., Boettiger, A.N., Moffitt, J.R., Wang, S., and Zhuang, X. (2015). RNA imaging. Spatially resolved, highly multiplexed RNA profiling in single cells. Science 348, aaa6090.

88. Koblan, L.W., Yost, K.E., Zheng, P., Colgan, W.N., Jones, M.G., Yang, D., Kumar, A., Sandhu, J., Schnell, A., Sun, D., et al. (2025). High-resolution spatial mapping of cell state and lineage dynamics in vivo with PEtracer. Science 390, eadx3800.

89. Soifer, H.S., Koch, T., Lai, J., Hansen, B., Hoeg, A., Oerum, H., and Stein, C.A. (2012). Silencing of gene expression by gymnotic delivery of antisense oligonucleotides. Methods Mol Biol 815, 333–346.

90. Stein, C.A., Hansen, J.B., Lai, J., Wu, S., Voskresenskiy, A., Høg, A., Worm, J., Hedtjärn, M., Souleimanian, N., Miller, P., et al. (2010). Efficient gene silencing by delivery of locked nucleic acid antisense oligonucleotides, unassisted by transfection reagents. Nucleic Acids Res 38, e3.

91. Bolger, A.M., Lohse, M., and Usadel, B. (2014). Trimmomatic: a flexible trimmer for Illumina sequence data. Bioinformatics 30, 2114–2120.

92. Dobin, A., Davis, C.A., Schlesinger, F., Drenkow, J., Zaleski, C., Jha, S., Batut, P., Chaisson, M., and Gingeras, T.R. (2013). STAR: ultrafast universal RNA-seq aligner. Bioinformatics 29, 15–21.

93. Shumate, A., Wong, B., Pertea, G., and Pertea, M. (2022). Improved transcriptome assembly using a hybrid of long and short reads with StringTie. PLoS Comput Biol 18, e1009730.

94. Markolin, P., Rätsch, G., and Kahles, A. (2022). Identification, Quantification, and Testing of Alternative Splicing Events from RNA-Seq Data Using SplAdder. Methods Mol Biol 2493, 167–193.

95. Liao, Y., Smyth, G.K., and Shi, W. (2014). featureCounts: an efficient general purpose program for assigning sequence reads to genomic features. Bioinformatics 30, 923–930.

96. Love, M.I., Huber, W., and Anders, S. (2014). Moderated estimation of fold change and dispersion for RNA-seq data with DESeq2. Genome Biol 15, 550.

97. Robinson, J.T., Thorvaldsdottir, H., Turner, D., and Mesirov, J.P. (2023). igv.js: an embeddable JavaScript implementation of the Integrative Genomics Viewer (IGV). Bioinformatics 39. 10.1093/bioinformatics/btac830.

98. Chu, C., Quinn, J., and Chang, H.Y. (2012). Chromatin isolation by RNA purification (ChIRP). J Vis Exp. 10.3791/3912.

99. Quinlan, A.R., and Hall, I.M. (2010). BEDTools: a flexible suite of utilities for comparing genomic features. Bioinformatics 26, 841–842.

100. Allan, M.F., Aruda, J., Plung, J.S., Grote, S.L., Martin des Taillades, Y.J., de Lajarte, A.A., Bathe, M., and Rouskin, S. (2024). Discovery and Quantification of Long-Range RNA Base Pairs in Coronavirus Genomes with SEARCH-MaP and SEISMIC-RNA. bioRxiv. 10.1101/2024.04.29.591762.

101. Darty, K., Denise, A., and Ponty, Y. (2009). VARNA: Interactive drawing and editing of the RNA secondary structure. Bioinformatics 25, 1974–1975.

102. Gagliardi, M., and Matarazzo, M.R. (2016). RIP: RNA Immunoprecipitation. Methods Mol Biol 1480, 73–86.

103. Abdulhay, N.J., Fiorini, C., Verboon, J.M., Ludwig, L.S., Ulirsch, J.C., Zieger, B., Lareau, C.A., Mi, X., Roy, A., Obeng, E.A., et al. (2019). Impaired human hematopoiesis due to a cryptic intronic splicing mutation. J Exp Med 216, 1050–1060.

